# *Dehalobacter* dechlorinates dichloroanilines and contributes to the natural attenuation of dichloronitrobenzenes at a complex industrial site

**DOI:** 10.1101/2025.02.13.638192

**Authors:** Sofia P. Araújo, Line Lomheim, Suzana P. Q. Kraus, E. Erin Mack, Jim C. Spain, Sávia Gavazza, Elizabeth A. Edwards

## Abstract

The potential for bioremediation of dichloroanilines (2,3- and 3,4-DCA) and dichloronitrobenzenes (3,4-, 2,5- and 2,3-DCNB) was investigated using inocula from an industrial site in northeast of Brazil. Anaerobic biotransformation of these chlorinated compounds was observed in microcosms simulating site conditions, particularly when electron donor was added. To disentangle specific transformation reactions, sub-cultures from active anaerobic microcosms were enriched with individual DCA or DCNB isomers. In these single-compound enrichment cultures, DCNB isomers were reduced to the corresponding DCA. Each DCA isomer was stoichiometrically dechlorinated to the corresponding monochloroaniline (CA) isomers. Further dechlorination of CA to aniline was noted. The reduction of DCNB to DCA was sustained by known fermenters and possibly *Desulfitobacterium*, while *Dehalobacter* was clearly responsible for dechlorination of DCAs. These findings helped to attribute function to microbial community data retrieved from field samples to inform the conceptual site model and enable identification of bioactive zones and rate-limiting conditions at a complex site. More generally, these data broaden the understanding of the metabolic repertoire of *Dehalobacter* and provide a model for incorporating microbial community data into site remediation efforts.

Graphical Abstract

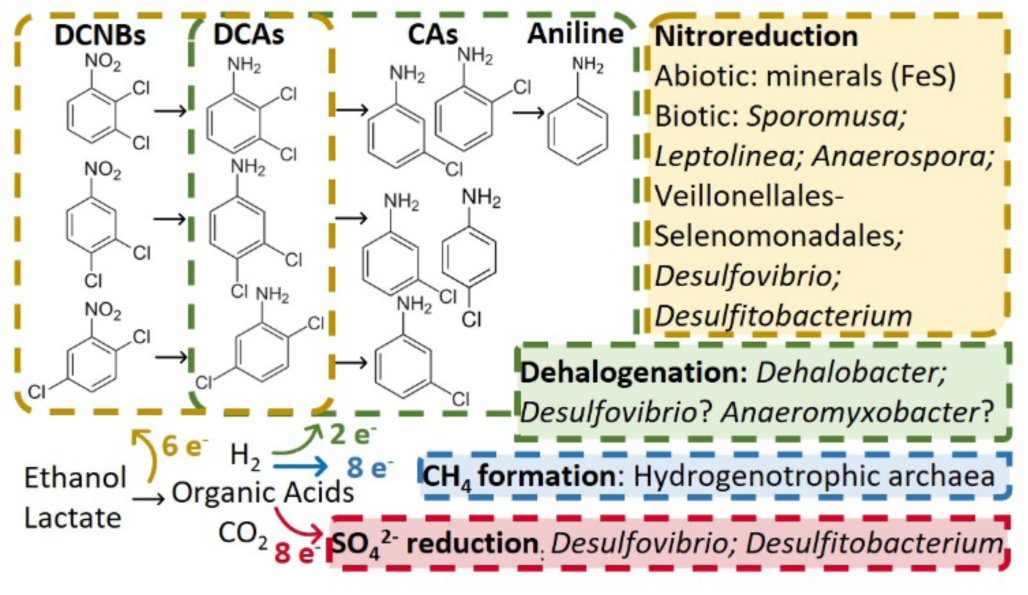

**Short synopsis statement:** A diverse microbial community contributes to natural attenuation at a complex contaminated site. Notably, dichloroanilines produced by biologically-mediated nitroreduction of dichloronitrobenzenes were dechlorinated by *Dehalobacter*.

## 1 INTRODUCTION

Many historically-contaminated sites worldwide are a challenge to remediate owing to complex mixtures of toxic contaminants and heterogeneous stratigraphy. For example, an industrial facility located in Bahia in NE Brazil has been the subject of several remedial investigations.^1,2^ As a former manufacturing facility of the herbicide diuron, it is impacted by high concentrations of multiple compounds including dichloronitrobenzenes (DCNB; ∼3,000 mg/kg) and dichloroanilines (DCA; ∼700 mg/kg) among many others.^1,2^ Anoxic conditions typically prevail in regions with low permeability, high organic matter, or high contaminant concentrations, where the rate of oxygen depletion exceeds rates of replenishment by diffusion or groundwater flow. Therefore, we sought to understand the fate of DCNB and DCA isomers under anoxic conditions with particular attention to associated microbial populations to elucidate microbial pathways and thus create a better conceptual model of the site and a more effective remediation strategy.

In subsurface anoxic environments, nitroaromatic compounds tend to become reduced to their corresponding aromatic amines. As early as 1975, the pesticide pentachloronitrobenzene was shown to be reduced to pentachloroaniline.^3^ Nitro group reduction to the corresponding amine is now understood to involve broad specificity and oxygen-independent nitroreductases and related enzymes that are widely distributed in bacteria and archaea.^4^ These reactions produce toxic transient nitroso and hydroxylamino intermediates, which are highly reactive and unstable.^4–6^ Nitroaromatic reduction may also be a respiratory process akin to nitrate reduction.^7–9^ A 1983 study reported the reduction of 3,4-DCNB to a variety of poorly quantified products including 3,4-DCA, and condensation products.^5^ Despite their environmental relevance, few studies have investigated the anaerobic biodegradation of DCNB isomers.

Literature on the anaerobic biodegradation of chloroanilines is also limited. The reductive dechlorination of 3,4-DCA to a mixture of monochloroaniline isomers (3-CA and 4-CA) under anaerobic/methanogenic conditions was first reported in 1989 and 1990, but the organisms responsible were not identified.^10–12^ Under nitrate-reducing conditions, a novel reductive deamination reaction was observed in a *Rhodococcus* strain where 3,4-DCA was deaminated primarily to dichlorobenzene,^13^ however, additional reports of this process could not be found. More recently, *Dehalococcoides mccartyi* strain CBDB1 and *Dehalobacter sp*. strain 14DCB1, both obligate organohalide-respiring-bacteria (OHRB) that dechlorinate chlorinated benzenes, were shown to dechlorinate 2,3-DCA to 2- and 3-CA and 3,4-DCA to 3-CA with no further transformation when acetate was provided as carbon source and hydrogen as electron donor.^14^

Reports of further dechlorination of monochloroanilines to aniline are sparse and inconclusive. Early reports (1997-98) described dechlorination of DCA to CA and then to small amounts of aniline under sulfate-reducing conditions.^15,16^ These studies did not characterize the microbial community. A 2019 study showed that *Geobacter* sp. KT5 could transform dichloroanilines under iron-reducing conditions; mineralization of the products via aniline and 4-aminobenzoate was proposed.^17^ Two years later, a mixed culture composed of *Geobacter* sp., *Paracoccus denitrificans*, *Pseudomonas* sp., and *Rhodococcus* sp. strains was described to transform propanil to 3,4-DCA, and further to 3-CA, chlorobenzene, and aniline, prior to mineralization.^18^ Another study found that 2-aminoanthraquinone-graphene oxide enhanced dechlorination in an enriched anaerobic sludge community that was able to dechlorinate 2-CA. In this work, some aniline and hexanoic acid were detected and organisms from *Oscillospira*, *Lactobacillales*, *Veillonellaceae*, and *Ruminococcus* groups were positively correlated with 2-CA transformation.^19^

Given the scarce literature on the biodegradation of dichloronitrobenzenes and dichloroanilines, the goal of this study was to identify organisms responsible for transformation reactions of such compounds and to use the information to design biomarkers to track biodegradation and delineate bioactive zones across a contaminated site. We determined that *Dehalobacter* plays a very active and critical role in the dechlorination of dichloroanilines at a complex site in Brazil. This knowledge is crucial for developing cost-effective *in situ* remediation strategies for many other complex contaminated sites.

## 2 MATERIALS AND METHODS

### 2.1 Chemicals and stock solutions

Substrates added to enrichment cultures in this study were 3,4-dichloronitrobenzene (3,4-DCNB), 2,5-dichloronitrobenzene (2,5-DCNB), 2,3-dichloronitrobenzene (2,3-DCNB), 3,4-dichloroaniline (3,4-DCA), and 2,3-dichloroaniline (2,3-DCA) (Sigma Aldrich, purity > 99%, (Figure S1). Stock solutions of each compound were prepared in acetone (Sigma Aldrich) at a concentration of 5 g/L. Ethanol (100%: Sigma Aldrich) and 60% sodium lactate (TCI America) were used as electron donors. The mentioned DCNB and DCA isomers, aniline, 2-, 3-, and 4-chloroaniline (CA), 2,5-DCA, DCB isomers (Sigma Aldrich, purity > 99%), and methane (Scott) were also used as analytical standards. Methanol (Sigma Aldrich) was used as the solvent for the chlorinated standards and acetonitrile (Fisher Scientific) was used for the mobile phase for liquid chromatography (described below).

### 2.2 Preparation of original microcosm study and enrichment cultures

Cultures were enriched from microcosms originating from a large laboratory treatability study done to evaluate the potential for in situ bioremediation of chlorinated and nitroaromatic compounds at an industrial site in Brazil. The original microcosms were constructed in 2015 with solid subsurface material and groundwater from the industrial site, incubated under anaerobic conditions at site pH (4.6 to 6.5), and amended with various electron donors and acceptors. The microcosms had mixtures of chloroanilines, dichloroanilines, dichloronitrobenzenes and chlorobenzenes at concentrations typical of the site, and biodegradation was monitored closely over a period of one year. After this, some of the most active microcosms continued to be monitored and re-amended with contaminant mixtures and electron donors (ethanol and lactate), and topped up with groundwater, as needed, until March 2020. At this time, six of these most active microcosms (microcosms Cam16, Cam17, Cam64, Cam65, Cam88, and Cam89), all established with electron donor to stimulate methanogenic conditions, were selected as inoculum for the bottles described in this study (Figure S1). The microcosm setup is detailed in Supporting Information Text S1 and in a Master’s Thesis.^20^ Results are summarized in Table S1 and Figure S2.

The first set of transfer cultures (34DCNB-T1, 25DCNB-T1, 23DCNB-T1, 34DCA-T1 and 23DCA-T1) and an autoclaved killed control (KC) were inoculated with cell pellets from one of the microcosms mentioned above (Figure S1). The pellet used as inoculum was obtained from a 40 ml sample from a shaken microcosm by centrifuging at 12,000 g for 15 min. These transfer cultures (group T1) were repeatedly amended with single individual chlorinated substrates: 3,4-DCNB, 2,5-DCNB, 2,3-DCNB, 3,4-DCA, and 2,3-DCA (Figure S1). The killed control (KC) was autoclaved three times on three consecutive days prior to being amended with a mix of six compounds: 3,4-DCNB, 2,5-DCNB, 3,4-DCA, 2,3-DCA, 2-CA, and aniline. An abiotic control (AC1) was also prepared, containing only reduced mineral medium supplemented with a slightly modified set of six compounds: 3,4-DCNB, 2,3-DCNB, 3,4-DCA, 2,3-DCA, 2-CA, and aniline. This modification was necessary because 2,5-DCNB and 2,3-DCNB co-elute under the chromatographic conditions used to measure compound concentrations.

A second generation of transfer cultures (34DCNB-T2, 25DCNB-T2, 23DCNB-T2, 34DCA-T2, and 23DCA-T2) was prepared from each T1 culture after ∼4 months. These were amended with the same individual compounds as the first generation transfers (3,4-DCNB, 2,5-DCNB, 2,3-DCNB, 3,4-DCA, and 2,3-DCA, respectively) (Figure S1). The DCNB-T2 cultures were inoculated with a 50% transfer from the corresponding T1 culture (50 mL into 50 mL fresh medium) while DCA-T2 cultures were inoculated with a 10% transfer from T1 cultures (10 mL of T1 into 90 mL). At the time we initiated the T2 cultures, the mechanism of DCNB reduction was unclear (discussed later). Therefore, we opted for a more conservative transfer approach, using a 50% inoculum instead of the 10% used for DCA cultures. Additional mineral medium supernatant was added to a T2 culture bottle when its volume was depleted to below 50 mL due to sampling, and the cultures were topped up to at least 80 mL. Occasionally, a little more medium was added to increase the volume of culture up to 250 mL. A second abiotic control (AC2) was prepared with mineral medium devoid of FeS and amended with 3,4-DCNB, 2,3-DCNB, 3,4-DCA, and 2,3-DCA. The data presented herein focus on the second transfer (T2) enrichment cultures.

All transfer cultures as well as controls were prepared in autoclaved 250-mL Boston round bottles sealed with Mininert caps, containing 100 mL of anaerobic mineral medium (pH 7) reduced with iron sulfide (FeS).^21^ Only medium supernatant was used to avoid adding precipitated FeS. To simulate site groundwater conditions and to promote mineralization of nitrogen-containing compounds, ammonium was omitted from the medium recipe. All bottles were incubated statically in the dark at room temperature in an anaerobic glovebox (Coy) supplied with a gas mix of N_2_:CO_2_:H_2_ (80:10:10%). No replicates were conducted.

The semi-volatile substrates (electron acceptors) were added as evaporated acetone stocks as described in Text S2. The target concentration was 10 mg/L (52 μM [DCNB] and 62 μM [DCA]. Second transfer enrichments had the concentration increased up to ∼27 mg/L (140 μM [DCNB] and 170 μM [DCA]) over time, to expose the cultures to a higher concentration of the corresponding compound for selection purposes. As electron donors, ethanol and lactate were added to the active cultures and the two controls (AC1 and KC) initially and were re-amended to active cultures when degradation rates slowed. Electron donors were added in ∼17-fold excess in electron equivalents (eeq) considering both nitroreduction and complete dechlorination of DCNB to aniline (Text S2). The initial pH was between 6.6 and 7.1 in all active enrichments.

### 2.3 Analytical procedures

Concentrations of DCNB, DCA and CA isomers and aniline were measured using a Hewlett-Packard 1050 series high-performance liquid chromatography (HPLC) system (Acclaim™ 120 C18 column, Thermo Scientific) equipped with a UV detector (set at 254 nm). The mobile phase was an isocratic 50:50 mixture of Milli-Q water and acetonitrile (Millipore Sigma) at 1 mL/min with a runtime of 25 min. Samples (2 mL) were taken from cultures using a 2 mL gastight glass syringe and filtered through a 0.22 μm (13 mm) Chromspec PTFE syringe filter (pre-wetted with methanol) prior to being injected into the HPLC. Lactate, acetate, propionate, formate, butyrate, pyruvate, chloride, nitrate, nitrite, sulfate, and phosphate were measured by ion chromatography (IC) using a Dionex^TM^ Integrion^TM^ IC system (Thermo Fisher Scientific). Methane was measured in headspace samples by gas chromatography (GC) using an Agilent 7890A gas chromatograph with headspace autosampler G1888 (Agilent Technologies). Method details are provided in Text S3.

### 2.4 Microbial community analysis and qPCR

Well-mixed samples (4 mL) from enrichment cultures were collected periodically and centrifuged at 10,000 rpm for 20 minutes. The supernatant was removed and cell pellets stored at −80°C. DNA was extracted using the DNeasy PowerSoil Kit (Qiagen) and quantified using a Qubit® 3.0 Fluorometer (Thermo Fisher Scientific). DNA samples were sent for Illumina MiSeq PE300 amplicon sequencing at an external laboratory (Genome Quebec Innovation Centre, Montreal, Canada), using modified “staggered end”^22^ versions of primers 926f-modified (5’-AAACTYAAAKGAATWGRCGG-3’) and 1392r-modified (5’-ACGGGCGGTGWGTRC-3’), which target the V6−V8 region of the 16S rRNA gene from both Bacteria and Archaea, as well as the 18S rRNA gene in Eukarya.^23,24^ Sequencing data was processed using QIIME 2 version 2021.2.^25^ After trimming the primer region with the cutadapt plug-in, amplicon sequence variants (ASVs) were generated using the DADA2 plugin,^26^ and taxonomy was assigned to the ASVs through the feature-classifier plugin using the SSU SILVA v.138 database.^27^ Data were further processed in Excel. Sequences <400 bp in length were removed and ASVs were ranked by maximum sequence abundance across all DNA samples. ASVs with abundance greater than 1% in at least one sample were kept for graphing. The Basic Local Alignment Search Tool (BLAST) available at the National Center for Biotechnology Information (NCBI) was used to compare ASV sequences of interest. To investigate the presence of the organisms of interest identified in the cultures, key genera were subsequently searched in a database containing 16S rRNA gene sequences retrieved from field samples collected by our collaborating partner during various remediation pilot projects conducted at the site. This database also contains all raw DNA sequences from microcosms and enrichments and was uploaded to the NCBI (Accession PRJNA1040917 as described in Table S2).

DNA samples were also analyzed by quantitative polymerase chain reaction (qPCR) using a CFX96TM real-time PCR detection system (Bio-Rad Laboratories Inc.). Primers GenBac1055f (5’-ATGGYTGTCGTCAGCT-3’) and GenBac1392r (5’-ACGGGCGGTGTGTAC-3’) targeting most bacteria^24,28,29^ and primers GenArch787f (5’-ATTAGATACCCGBGTAGTCC-3’) and GenArch1059r (5’-GCCATGCACCWCCTCT-3’) targeting most archaea^30^ were used. A no-template control was also included in each run. All qPCR samples were run in triplicate. Details on the qPCR method are provided in Text S4. Absolute total bacterial and archaeal abundance obtained by qPCR (copies 16S rRNA gene per ml) was used to convert the relative abundances obtained from amplicon sequencing to absolute abundance for each ASV.

A total of forty-eight DNA samples were sequenced and analyzed by qPCR, including samples from the original subsurface solid material, microcosms, and enrichments. On average 66,000 16S rRNA gene amplicon reads were retrieved per sample (Table S3a). Reads were grouped by bacteria and archaea, then sequences with relative abundance >1% in any sample (Table S3b for bacteria and Table S3c for archaea) were used to calculate absolute abundances (copies/mL) of each ASV using total abundances of bacteria and archaea determined by qPCR (Table S4). Final absolute abundances for each ASV (copies/mL) are reported in Tables S5a to S5f for bacteria; Tables S6a to S6f for archaea. Finally, Non-metric Multi-dimensional Scaling (NMDS) and bubble plots were constructed as explained in Text S5.

## 3 RESULTS AND DISCUSSION

### 3.1 Abiotic transformation of DCNB was not sustained

Contaminant profiles in three controls (AC1, AC2 and KC) were assessed for possible abiotic reactions. Chloroanilines and aniline were not transformed in any of these controls. Dichloronitrobenzenes were not transformed in medium control (AC2) without FeS (Figure S3; Table S7). In the medium control (AC1) with FeS, 3,4-DCNB was not transformed, but 2,3-DCNB was reduced to 2,3-DCA (Figure S4; Table S7); however, this activity was not sustained upon reamending with 2,3-DCNB. These data are consistent with reports that reduced minerals, such as FeS or perhaps reduced resazurin, can sometimes mediate reduction of the nitro group. ^31–38^ Curiously, DCNB isomers were repeatedly transformed to DCA in the autoclaved control KC (Figure S5; Table S7); however later analyses (to be discussed) revealed that KC was in fact not sterile.

### 3.2 Biologically-mediated nitroreduction and dechlorination

Each transfer culture established on single substrates remained active, sustained by refeeding chlorinated substrates (DCNB or DCA) and electron donors as needed for over 500 days. Historical data are provided in Table S8 (T1) and Table S9 (T2). Focusing on T2 transfers from day 280 to 520 and on the three DCNB-amended cultures first, we observed consistent and complete transformation of each DCNB isomer to the corresponding dichloroaniline (Figure 1a-c). In the culture amended with 3,4-DCNB the product, 3,4-DCA, was further dechlorinated to 3- and 4-CA (Figure 1a), while in the cultures amended with 2,3- and 2,5-DCNB, the transformation stopped at the corresponding dichloroaniline (Figures 1b and 1c). The sum of measured chlorinated products agreed well with the cumulative DCNB added. Reduction of 2,3-DCNB (<∼1 µM per day) was ∼5 times slower than the reduction of either 3,4- or 2,5-DCNB (5-6 µM per day).

**Figure 1.**
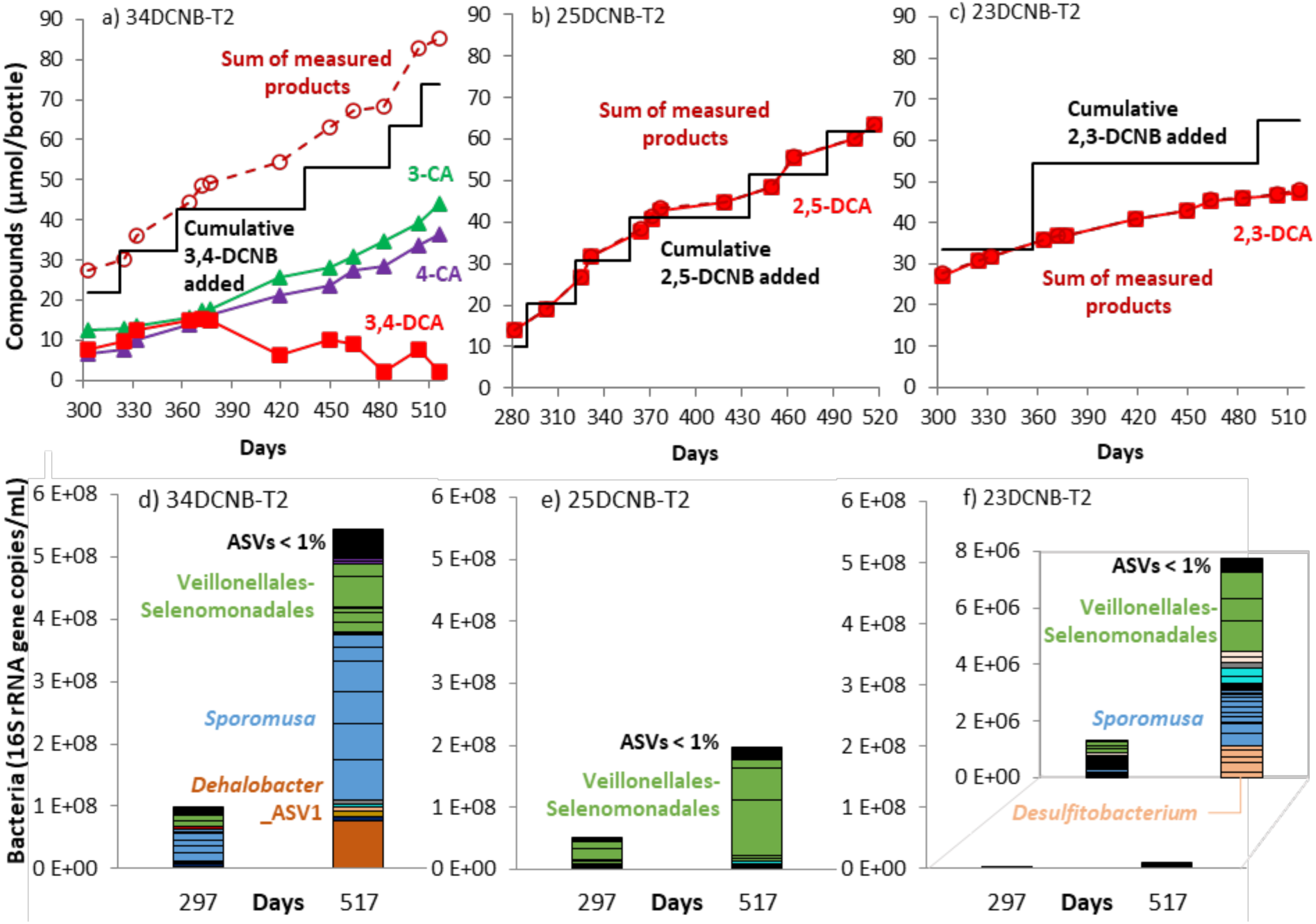
Degradation profiles (upper panels a), b) and c); µmole/bottle) and corresponding major microbial taxa (lower panels d), e) and f); copies/mL) in DCNB-amended T2 transfer cultures from day 280 to 520. Cumulative DCNB added (––––) is shown for each amendment (step) of 70 to 140 µM (10 to 20 µmol per bottle). The sum of products 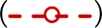, including DCA 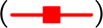, 4-CA 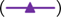, 3-CA 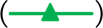, 2-CA and aniline (not shown, mostly below detection), illustrates a reasonable mole balance. Measured DCNB concentrations are not shown for clarity but can be found in Table S9. In lower panels amplicon sequence variants (ASVs) from the same taxonomic designation are assigned the same colour.

Focusing next on the two DCA-amended transfers, 3,4-DCA was dechlorinated to a mix of 3- and 4-CA (Figure 2a), while 2,3-DCA was dechlorinated primarily to 3-CA, with traces of 2-CA (Figure 2b). These cultures dechlorinated about ∼5 to 6 µM DCA to CA per day.

**Figure 2.**
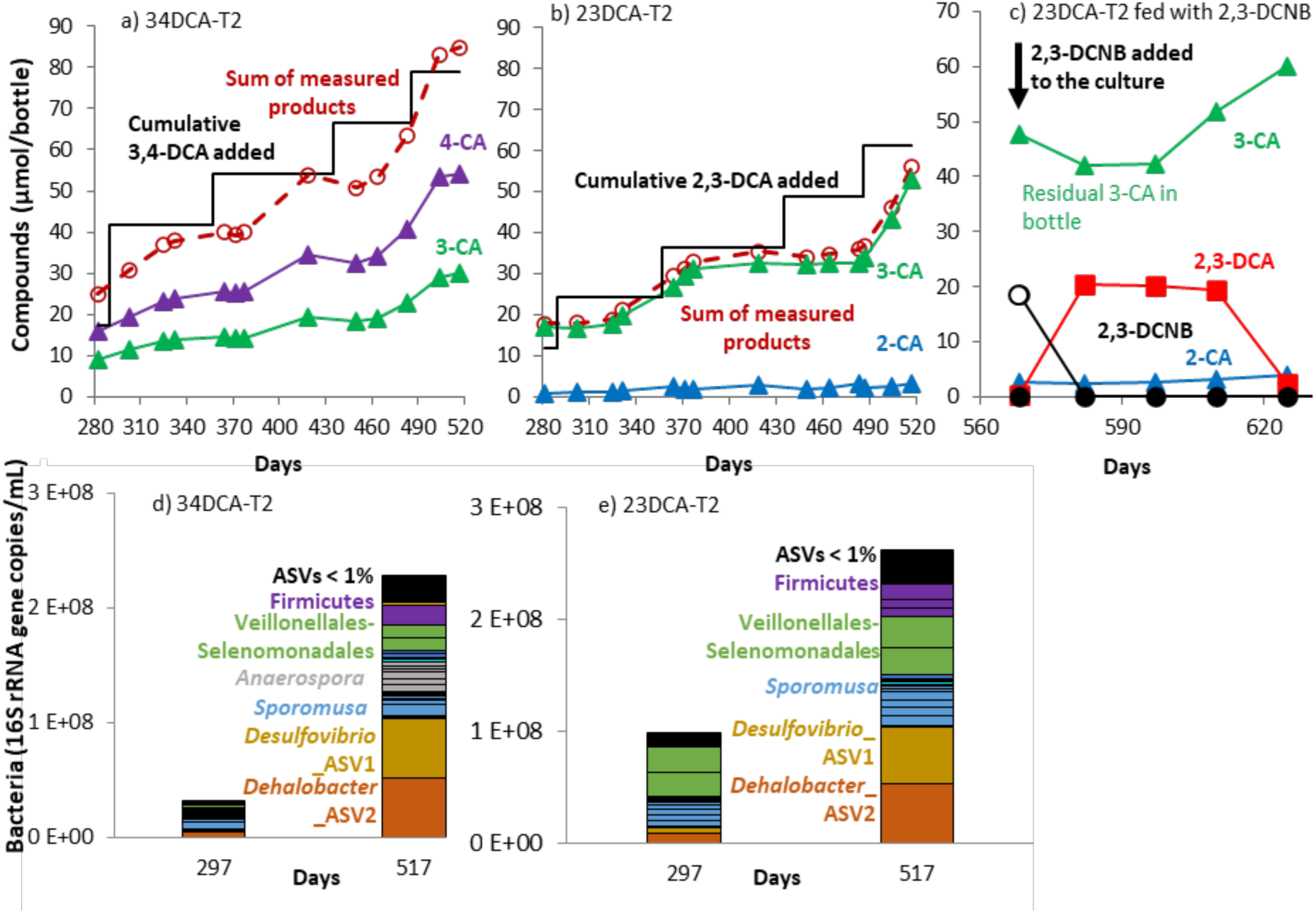
Degradation profiles (upper panels a), b) and c); µmole/bottle) and corresponding major microbial taxa (lower panels, d) and e); copies/mL) in two DCA-amended T2 transfer cultures from day 280 to 520 (panels a), b), d), and e)) and from day 560 to 625 (panel c)). In panels a) and b), cumulative DCA added (–––) is shown for each amendment (step) of ∼80 µM (∼12 µmol per bottle). The sum of products 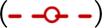, including DCA 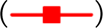, 4-CA 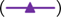, 3-CA 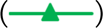, 2-CA 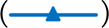, and aniline (not shown, mostly below detection), illustrates a reasonable mole balance. Measured DCA concentrations are not shown for clarity but can be found in Table S9. Panel c) shows degradation profile after 2,3-DCA-T2 was fed with 2,3-DCNB 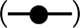 instead of 2,3-DCA on Day 568. The open circle shows the amount of 2,3-DCNB fed to 23DCA-T2 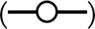. In lower panels amplicon sequence variants (ASVs) from the same taxonomic designation are assigned the same color.

Given the very slow transformation rate in 23DCNB-T2, we wondered if the enrichment culture dechlorinating 2,3-DCA (23DCA-T2; Figure 2b) could also reduce 2,3-DCNB, even though this culture - and the microcosm (Cam16) from which it was derived - had never been exposed to any DCNB isomer. Surprisingly, 2,3-DCNB was reduced without a lag to 2,3-DCA which was further dechlorinated to 3-CA (Figure 2c). We then tested all active DCA-degrading cultures with a mix of the three DCNB isomers; all the cultures were able to reduce DCNBs to their respective DCA isomer (data not shown).

### 3.3 Microbial community composition reflects electron balance

To relate the microbial community to activity, we first identified abundant bacterial and archaeal ASVs (>1%) and then calculated their absolute abundance in active cultures focusing on 2-time points for each sample. Figure 3 provides a summary of the most relevant taxa identified. Most of the enriched bacterial ASVs belong to the phylum Bacillota (Firmicutes) including *Sporomusa*, *Anaerospora*, and unassigned members of the Veillonellales-Selenomonadales. These organisms clearly grew on the ample electron donor amendments to the cultures. ASVs for *Anaeromyxobacter*, *Desulfitobacterium* and *Desulfovibrio* were also retrieved. These genera are also often implicated in nitrate or sulfate reduction though some members may be facultative dechlorinators. ASVs for obligate dechlorinators of the genus *Dehalobacter* were also identified (Figure 3). A complete overview of all major taxa (>1%) are shown in Figure S6, for bacteria, and Figure S7, for archaea.

**Figure 3.**
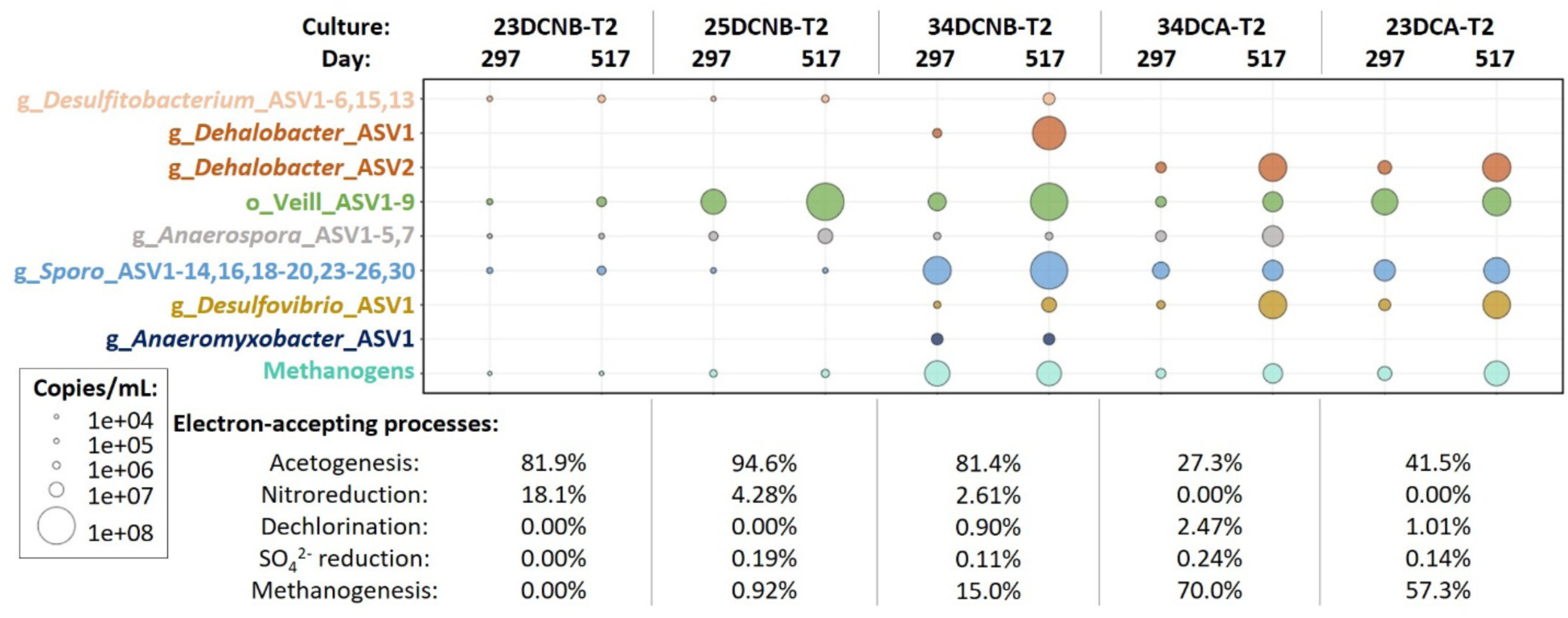
Absolute abundance of major organisms (copies/mL) and proportion of electron equivalents from amended donors channeled to each electron-accepting processes (% electron equivalents) for days 297 and 517. Veillonellales-Selenomonadales order is shortened to “Veill”; *Sporomusa* genus is shortened to “*Sporo*”. Calculated absolute abundances for all ASVs are available in Table S5 (bacteria) and Table S6 (archaea).

To confirm the extent of dechlorination, an electron balance of possible donors and acceptors was completed using measured concentrations of chlorinated organics, methane (Table S9) and anions (Table S10). As expected from stoichiometry (e.g., C_6_H_5_Cl_2_N + H_2_ → C_6_H_6_ClN + HCl), chloride concentrations increased proportionally in cultures that exhibited dechlorination (34DCNB-T2, 34DCA-T2, and 23DCA-T2). The measured increase in chloride was slightly higher than expected in almost all cases (Table S11), likely because of the error associated with measuring small changes against a high background of ∼1900 μM chloride.

Potential electron-accepting processes include acetogenesis, methanogenesis, nitroreduction, dechlorination, and sulfate reduction. For T2 cultures between days 280 and 520, we calculated the proportion of electron equivalents from added donors channeled to each of the different electron-accepting processes (Figure 3, bottom table). In DCNB-amended cultures, acetogenesis from lactate and ethanol was the dominant process, accounting for over 80% of electrons from donors. Methanogenesis was severely limited. By contrast, in DCA-amended cultures, methanogenesis accounted for 60-70% of electron equivalents from donors, which combined with acetogenesis accounted for over 98% of electron equivalents. A buildup of acetate in all cultures (Tables S10 and S12) suggested possible inhibition of acetoclastic methanogens specifically, particularly by DCNBs. Culture 34DCNB-T2 produced the most methane among DCNB-amended cultures but also exhibited the most rapid and extensive nitroreduction and dechlorination, thus relieving inhibition associated with DCNBs and DCAs.

Other electron accepting processes accounted for a much lower proportion of activity (Figure 3). Nitroreduction consumed 5% of electron equivalents in cultures fed with 3,4- and 2,5-DCNB isomers and up to 20% of electron equivalents in culture 23DCNB-T2, although activity in this culture was very low and no lactate was consumed (Tables S10 and S12). In all cases, dechlorination accounted for less than 3% of all electron equivalents provided. The small amount of sulfate (∼30 µM) present in the medium was rapidly depleted in the 4 highly active cultures (34DCNB-T2, 25DCNB-T2, 34DCA-T2, and 23DCA-T2). However, less than 1% of electron equivalents were channeled to sulfate reduction. In culture 23DCNB-T2, sulfate was not consumed, consistent with low activity observed in this enrichment (Table S11).

We compared the proportions of electrons consumed by each electron-accepting process with corresponding microbial proportions (Figure 3) to infer function (Table S12). Methane production was greatest in cultures 23DCA-T2 and 34DCA-T2, lower in 34DCNB-T2, very low in 25DCNB-T2 and absent in 23DCNB-T2. Methanogens were correspondingly most abundant in 23DCA-T2, 34DCA-T2, and 34DCNB-T2. ASVs corresponding to hydrogenotrophic *Methanoregula* dominated the archaeal population, with a much lower abundance of *Methanocella* and other hydrogenotrophic genera (Figure S7). The presence of only hydrogenotrophic methanogens combined with substantial accumulation of acetate strongly indicates that DCNB isomers and to a lesser extent DCA isomers inhibited acetoclastic methanogens.

### 3.4 Nitro group reduction can be mediated by many different microbes

We had hoped to observe enrichment of specific microbes in DCNB-amended cultures to enable functional attribution. Rather, it seems that many different genera can likely reduce the nitro group. Biologically regenerated FeS is unlikely to play a role as a reductant, as there was no evidence of sulfate or iron cycling in the cultures, and the rates of DCNB reduction were similar regardless of FeS concentrations. In all cultures, there was significant representation of fermenting or acetogenic microbes, particularly Veillonellales-Selenomonadales, *Sporomusaceae*, *Sporomusa*, and *Anaerospora* (Figure S6). This is consistent with the ample supply of ethanol and lactate to all cultures.

We wondered if these fermenting and acetogenic organisms could also be reducing DCNB to DCA as a growth-associated electron-accepting process akin to nitrate reduction. In 25DCNB-T2, where only nitroreduction was observed without dechlorination, 6 ASVs (ASV1 to 5 and 8) (>90% identity) classified only to the order level as Veillonellales-Selenomonadales were predominant (Table S5b). They represented more than 80% of the bacterial community in this culture on day 517 (Figure 1e). The closest matches to these sequences are affiliated with the genera *Pelosinus*, *Anaerospora*, *Sporomusa*, *Dendrosporobacter*, among other organisms; interestingly, one of the closely matching sequences (KF460373.1) comes from a study on the impact of nitrate on pentachlorophenol dechlorination^39^ (Figure S8).

In fact, ASVs from Veillonellales-Selenomonadales were abundant in all 5 active cultures (Figures 1d, e, and f, Figures 2d and e and Figure 3), whether fed DCNB or DCA, making functional attribution difficult. Finding that DCNB isomers could be reduced in all cultures regardless of enrichment substrate is perhaps then not so surprising. An autoclaved control (KC) amended with a mixture of compounds (3,4-DCNB, 2,5-DCNB, 3,4-DCA, 2,3-DCA, 2-CA, aniline) was prepared to try to distinguish between biotic and abiotic reactions. Unfortunately, the bottle did not achieve sterility, despite autoclaving it on three consecutive days. In this “killed” control, both 3,4-DCNB and 2,5-DCNB were reduced to the corresponding DCA isomers (Figure S5). DNA was readily extracted from this control, and although the total microbial abundance was an order of magnitude lower than other cultures, the KC culture was dominated by a unique Butyricoccaceaea ASV as well as other spore-forming and presumably heat-resistant microbes. Among the microbes detected in KC after refeeding 3,4- and 2,5-DCNB 4 times, *Desulfitobacterium* and *Anaerospora* were also detected in other active nitroreducing cultures (i.e., 23DCNB-T2 and 25DCNB-T2) (Figure S6), suggesting these genera are also candidates for nitroreduction. Ultimately it became clear that nitroreduction is a widespread microbial activity that was easy to stimulate with electron donor in the cultures, however, the contribution of biotic nitroreduction to microbial growth is yet to be investigated.

### 3.5 Dechlorination of 2,3- and 3,4-DCA supports growth of *Dehalobacter*

In actively dechlorinating cultures, *Dehalobacter* was the only microbe to systematically show a substantial increase in abundance. *Dehalobacter* was not detected in cultures that were only catalyzing nitroreduction (25DCNB-T2 and 23DCNB-T2). Given that it is an obligate organohalide respiring bacterium, we conclude that *Dehalobacter* was responsible for the DCA-dechlorinating activity in enrichment cultures amended with 3,4-DCNB, 3,4-DCA, and 2,3-DCA (Figures 1d and 2d and e).

Two distinct *Dehalobacter* ASVs were identified: ASV1 in 34DCNB-T2 and ASV2 in 34DCA-T2 and 23DCA-T2 (Text S6), with maximum bacterial abundances of 14, 25, and 31% in each culture, respectively (Table S3b). ASV1 (468 bp) is 100% identical to several previously published 16S rRNA gene sequences of *Dehalobacter restrictus* strains, including strain SAD (CP148032.1), strain DSM 9455 (CP007033.1), and strain 12DCA (CP046996.1), as well as to three *Dehalobacter* strains classified only to the genus level, including strain DCA (CP003869.1), strain DAD (CP148031.1), and strain CF (1CP003870.1). The known substrates for these *Dehalobacter* strains include tetra- and trichloroethene,^40^ 1,2-dichloroethane,^41^ tri- and dichloromethane,^42^ and 1,1,1-trichloroethane and 1,1-dichloroethane.^43,44^

*Dehalobacter* ASV2 (468 bp) was not a perfect match to any 16S rRNA gene sequence in the NCBI database. It shares only 97.01% identity to ASV1 and 98.93% to *Dehalobacter* strain DCM (CP092282.1) described as “dichloromethane-degrading bacterium from Xi river” (Direct Submission by Yang, Y. *et al*., 13-Jan-2022). Neither *Dehalobacter* ASV1 nor ASV2 sequences matched (<95% identity) to the 16S rRNA gene fragment (937 bp) from *Dehalobacter* sp. strain 14DCB1 (JN051267.1), a strain enriched on dichlorobenzene^45^ but reported to dechlorinate dichloroaniline.^14^

To further confirm the role of *Dehalobacter* in chloroaniline dechlorination, we estimated growth yields from qPCR and concentration data to compare with previously published yields. The net growth yields ranged from 1.4 × 10^13^ to 6.5 × 10^13^ cells per mol of chloride released assuming 3 copies of the 16S rRNA gene per cell (Table S13). The yields calculated in this study are consistent with many previously reported yields for other *Dehalobacter* strains using various chloroethanes, chlorobenzenes, and chloromethanes as electron acceptors for growth (Table S14). In fact the yields align with the highest yields previously reported, solidifying chloroaniline dechlorination as a growth-related process.

Additional evidence for *Dehalobacter* being responsible for DCA dechlorination was obtained when non-dechlorinating culture 25DCNB-T2 was inoculated with 3 mL of dechlorinating culture 34DCNB-T2 (on day 653). After inoculation, residual 2,5-DCA previously formed from 2,5-DCNB was dechlorinated to 3-CA and the relative abundance of *Dehalobacter*_ASV1 increased from zero on day 517 to ∼9% on day 653 (Figure S9).

A shift in activity in culture 23DCA-T2 was noted as monitoring continued for over 1000 days (Figure S10). As time progressed, 2-CA rather than 3-CA became the major product from 2,3-DCA. Thereafter, 2-CA was slowly yet consistently transformed to aniline. On day 881, *Dehalobacter*_ASV2 constituted 16% of the bacterial community (Table S5e). 2-CA was dechlorinated to aniline in subsequent transfers from this culture as well (data not shown). A subsample of 23DCA-T2 was taken on day 653 for additional growth experiments to ultimately build up enough culture to measure compound specific isotopic enrichment factors. This ongoing research implicates *Dehalobacter* in aniline formation as described in a companion publication.^46^

### 3.7 Other organisms identified in enriched cultures

Several other ASVs belonging to known facultative OHRB were identified, including *Anaeromyxobacter*, *Desulfovibrio*, and *Desulfitobacterium* (Figure 3). While some members of these genera can dechlorinate, most are known to be quite versatile and may reduce sulfate, nitrate or even grow fermentatively in the absence of a terminal electron acceptor. An *Anaeromyxobacter* ASV found in culture 34DCNB-T2 is 99.79% similar over ∼450 bp to the 16S rRNA gene of *Anaeromyxobacter* sp. strain RF71 (MZ254878.1), 99.57% similar to *A. dehalogenans* 2CP-C (CP000251.1) and 99.36% similar to *A. dehalogenans* 2CP-1 (CP001359.1). Both strains 2CP-C and 2CP-1 can gain energy from the reductive dechlorination of chlorophenol and contain putative dehalogenases.^47–49^ However, the abundance of *Anaeromyxobacter* in the culture was far less than *Dehalobacter* and is therefore unlikely to be primarily responsible for dechlorination.

A single ASV of *Desulfovibrio* increased in abundance over time in the three cultures dechlorinating DCA isomers (Figure 3). *Desulfovibrio* are sulfate-reducing bacteria that can also ferment substrates such as lactate; some can respire organohalides.^50,51^ In cultures 23DCA-T2 and 34DCA-T2, *Desulfovibrio* increased between days 280 and 520 to similar extent as *Dehalobacter* (Figures 2d and 2e). However, a similar increase was not seen in dechlorinating culture 34DCNB-T2 (Figure 1d). The amount of sulfate in the medium (around 30 μM) is too small to explain this growth, therefore *Desulfovibrio* was more likely metabolizing lactate to acetate, possibly participating in dechlorination alongside *Dehalobacter*. The *Desulfovibrio* ASV in our cultures is 100% identical to a non-dechlorinating *Desulfovibrio* sp. (JQ599691.1) detected in a dichloroethane dechlorinating enrichment.^52^ The sequence was also 100% identical to an uncultured *Desulfovibrio* clone (AB854345.1) found in a non-dechlorinating environment.^53^ Therefore, the more likely role of this organism is fermentation and acetogenesis, in support of dechlorinating *Dehalobacter*.

Several ASVs (sharing > 94% identity) were assigned to *Desulfitobacterium*. Interestingly, these ASVs were only found in the cultures reducing DCNBs and in the KC control that exhibited nitroreduction; they were not detected in the DCA-amended cultures (Figure 3). This seems to implicate *Desulfitobacterium* in nitroreduction, however the ASVs were in low abundance. Furthermore, DCA-amended cultures could also rapidly reduce the nitro group, therefore *Desulfitobacterium* was not solely responsible for nitroreduction, if at all. Moreover, the 8 most abundant *Desulfitobacterium* ASV sequences are > 95% similar to a *Desulfitobacterium* sequence found in *Dehalococcoide*s-containing dechlorinating cultures (AB596883.1) (Figure S11), where they were not implicated in dechlorination,^54^ a likely scenario in our study as well.

### 3.8 Data integration and application to improving remediation outcomes

Non-metric multidimensional scaling (NMDS) was used to evaluate relationships among cultures as they were enriched from subsurface material to initial microcosms and finally to enrichment cultures (Figure 4). First, there is clearly an impact of the original source material, shown in red, blue and green for three original sediment sources used to construct the microcosms. Second, there was a convergence of community depending on substrate. Enrichment cultures that dechlorinate DCAs are distinct from those that only carry out nitroreduction. Samples from 34DCNB-T2 culture, in which both nitroreduction and dehalogenation were observed, are located between the dechlorinating-exclusive culture samples (34DCA-T2, and 23DCA-T2) and the nitroreducing-exclusive cultures (25DCNB-T2, 23DCNB-T2 and KC). *Dehalobacter* in the top groupings and *Desulfitobacterium* in the bottom two cultures (23DCNB-T2 and KC) are major genera driving the vertical spread of samples in the NMDS plot. Relationships between original microcosms and transfers are also illustrated in Figure S1.

**Figure 4.**
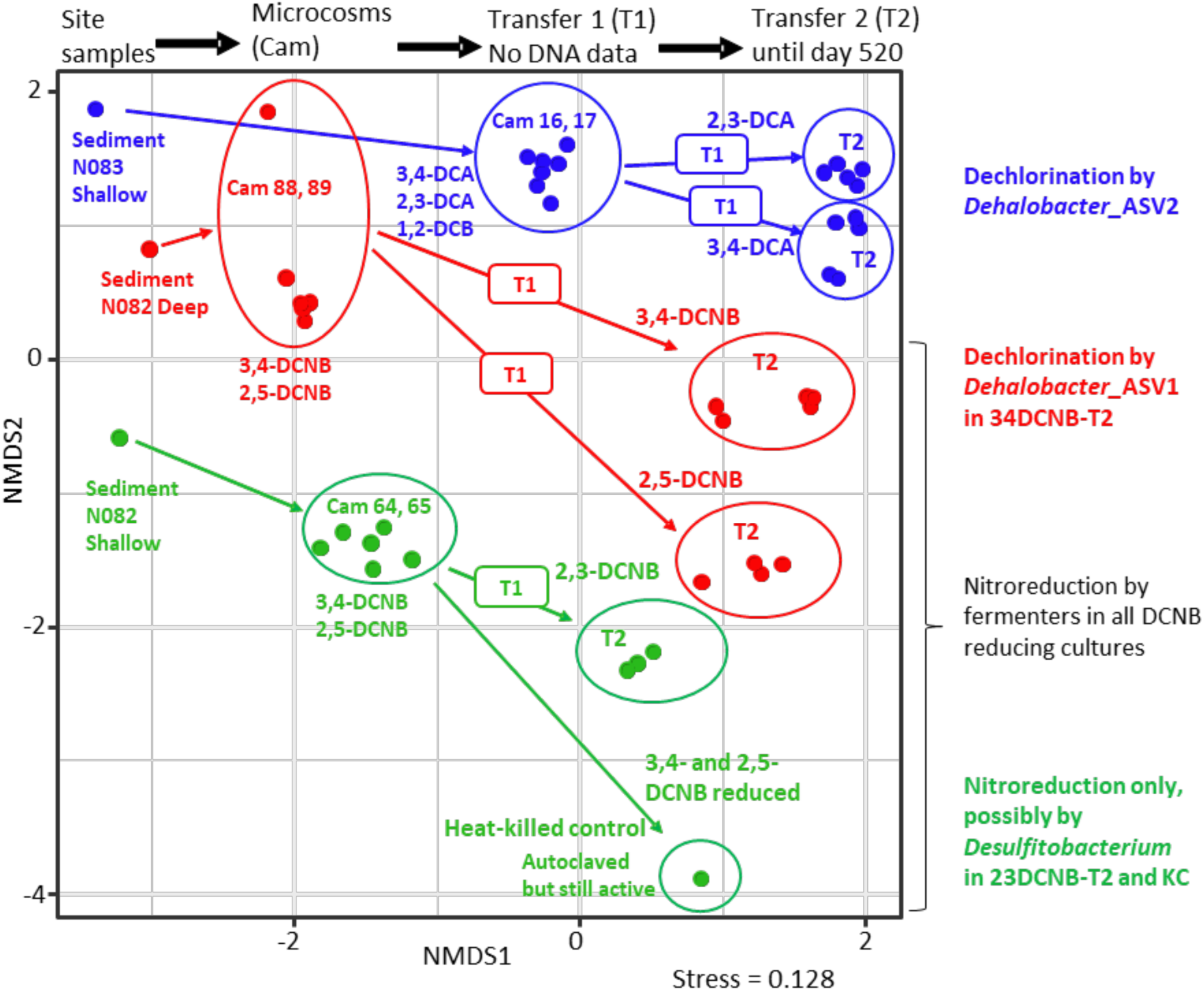
Non-metric multidimensional scaling plot based on relative abundance of 16S rRNA amplicons in 45 samples from the original site material used for microcosms and enrichment cultures.

In a survey of over 60 groundwater samples from around the site in Brazil that were analyzed by amplicon sequencing, approximately 14% returned *Dehalobacter*. Of the ∼400 samples that have been sequenced from various remediation trials implemented at the industrial site – including samples from pilot wetlands and pre and post biosparging units – many different *Dehalobacter* ASVs (including those enriched in this study) were detected in 78 samples (∼20%) clustered around more impacted zones with lower redox potential. Relative abundances of *Dehalobacter* reached 2% of the bacterial population in some locations, which is high considering the complex mixture of chemicals and conditions at the site.^1,2^ These abundances are being used to delineate biologically active zones and to predict attenuation rates to most effectively remediate the site.

Other ASVs detected in the enrichment cultures were detected in site samples, including the ones affiliated with *Anaeromyxobacter* and *Desulfitobacterium*, which were detected in ∼53% and ∼18% of the total ∼460 samples collected by the partner company, respectively. The ASVs classified as Veillonellales-Selenomonadales were detected in ∼11% of the same group of samples, while the ones classified further to *Sporomusa* and *Anaerospora* genera in ∼3% and ∼6%. The high relative abundance of the Veillonellales-Selenomonadales order in our enriched cultures indicates those organisms were favoured by the ample availability of donor (ethanol and lactate) added to the cultures, which is not the case in the field.

The fact that the microbes identified in enrichment cultures are also present at the site attests to the utility of such enrichments in site characterization and provides very strong evidence that these processes are contributing to active natural attenuation in anaerobic zones. At the site, concentrations of dichloroanilines are generally higher than dichloronitrobenzenes, consistent with our finding that biologically-mediated nitroreduction occurs readily and that there are many genera capable of catalyzing this reaction. It is highly likely that some of the other contaminants at the site could be electron donors, including petroleum hydrocarbons, solvents and plasticizers. In addition, abiotic reactions may also occur driven by reduced minerals or by reduced natural organic matter, which can donate electrons for nitroreduction.^37,55–58^ Further studies will seek to better understand the players and nitroreduction mechanisms in DCNB cultures.

Two distinct *Dehalobacter* ASVs were unequivocally linked to dechlorination, and their presence along with other ASVs in site samples is indicative of dechlorinating activity in situ. The two ASVs originated from geographically distinct locations at the site. Traces of aniline were found and continue to accumulate in 23DCA-T2, suggesting further dechlorination of CA is possible in situ. Inhibition of acetoclastic methanogens was noted, which is also consistent with site data and provides a means to evaluate site recovery: as DCNB and DCA concentrations decrease with active remediation efforts, methanogenic populations in anoxic zones should rebound. Biomarkers based on 16S rRNA genes are already being deployed at the site to track population changes as remediation alternatives are evaluated. The robust enrichment cultures generated herein will be invaluable for future studies of biotransformation mechanisms, development of isotopic enrichment factors and of additional biomarkers, and possibly even for use as bioaugmentation cultures.

## SUPPORTING INFORMATION

The following files are available free of charge:

Additional details on experiments, materials and methods, and data obtained, including texts and figures mentioned in this paper (PDF); Tables including data obtained and calculations mentioned in this paper (.xls).

## ACKNOWLEDGEMENTS

For the financial support for Sofia Araújo, we thank FACEPE (Science and Technology Foundation of the State of Pernambuco, Brazil – process number BFD-0013-3.07/21), CNPq (National Council for Scientific and Technological Development, Brazil - process number 165130/2017-2), and CAPES (Coordination for the Improvement of Higher Education Personnel) for the Graduate Program support (Proap) and Institutional Internationalization Program (PrInt) (process number 88887.363316/2019-00). This work was also supported by Mitacs through the Mitacs Accelerate and Mitacs Elevate programs (Application ref. IT23315 and IT34875, Canada).

The authors also acknowledge the grant from CNPq (process number 304862/2018-5) and FACEPE (process number APQ 0456-3.07/20), and NSERC (Natural Sciences and Engineering Research Council of Canada) Alliance project (ALLRP 575243-22), which has Geosyntec Consultants and Corteva Agriscience™ as the two main industrial partners.

The authors do not have a conflict of interest to declare. This publication is in accordance with the Brazilian biodiversity law n.° 13.123, from May 20^th^, 2015. This is non-profit research and has no commercial intention between the parties. This is a collaboration project between the parties involved. Access Registration No. A3D9F11.

We would also like to thank Elodie Passeport, Shuping Wang, and Luiz Pereira for their helpful discussions and collaborative work on this project.

