## Supplemental Information for "*Dehalobacter* dechlorinates dichloroanilines and contributes to the natural attenuation of dichloronitrobenzenes at a complex industrial site"

**Table of contents**


| Material | | Page |
|---|---|---|
| Text S1 | Original microcosm study setup and results. | S5 |
| Text S2 | Amendments to enrichment cultures and controls. | S6 |
| Text S3 | Details of analytical procedures (pH, HPLC, IC, GC). | S6 |
| Text S4 | Quantitative polymerase chain reaction (qPCR). | S9 |
| Text S5 | Preparation of Bubble Plots and Non-metric multidimensional Scaling (NMDS) analysis. | S9 |
| Text S6 | 16S rRNA gene sequences of *Dehalobacter* ASV1 (detected in 34DCNB-T2) and *Dehalobacter* ASV2 (detected in 23DCA-T2 and 34DCA-T2) obtained from 16S Illumina sequencing. | S9 |
| Figure S1 | History of enrichment cultures, showing setup details of original microcosms, T1, and T2 transfers. | S11 |
| Figure S2 | Overview of observed transformations deduced in active anaerobic microcosms. | S12 |
| Figure S3 | Concentration profiles in control bottle AC2, medium with no FeS. | S13 |
| Figure S4 | Concentration profiles in control bottle AC1, medium with FeS. | S14 |
| Figure S5 | Concentration profiles in autoclaved (killed) control bottle KC. | S15 |
| Figure S6 | Overview of the most abundant bacterial ASVs in the active bottles (ASVs > 1%, 2 time points), and activities observed in bottles. | S17 |
| Figure S7 | Overview of the most abundant archaeal ASVs in the active bottles (ASVs > 1%, 2 time points), and activities observed in the bottles. | S17 |
| Figure S8 | Phylogenetic tree of organisms classified as part of the Veillonellales-Selenomonadales order. | S18 |
| Figure S9 | Activity in 25DCNB-T2 between days 500 and 920, before and after the culture was inoculated with 3 mL of 34DCNB-T2 at T=653 (a). DNA samples collected from 25DCNB-T2 at T=517 and T=881, showing growth of *Dehalobacter*_ASV1 (same ASV | S19 |

| | | |
|---|---|---|
| | detected in 34DCNB-T2 went from 0 to ~10% of the bacterial community) (b). | |
| Figure S10 | Shift in activity in culture 23DCA-T2: 2-CA production and progressive accumulation of aniline. | S20 |
| Figure S11 | Phylogenetic tree of organisms classified as part of the *Desulfitobacterium* genus. | S21 |
| Table S1 | Summary of setup and activity observed in original anaerobic microcosms. | Excel |
| Table S2 | DNA samples described in this paper along with their corresponding names in the NCBI bioproject. | Excel |
| Table S3a | Results of 16S Illumina sequencing from subsurface material, original microcosms, and enrichment cultures, as raw reads and including sequences. | Excel |
| Table S3b | Results of 16S Illumina sequencing from subsurface material, original microcosms, and enrichment cultures, calculated as percent of bacterial reads, based on Table S3a. | Excel |
| Table S3c | Results of 16S Illumina sequencing from subsurface material, original microcosms, and enrichment cultures, calculated as percent of archaeal reads, based on Table S3a. | Excel |
| Table S4 | Concentrations of general bacteria and general archaea in enrichment cultures transfer 2 (T2) and heat-killed control (KC) as measured by qPCR of 16S rRNA genes (including calibration curves). | Excel |
| Table S5 | Absolute abundance of bacterial ASVs, calculated based on the Illumina sequencing data and on qPCR: a) 34DCNB-T2, b) 25DCNB-T2, c) 23DCNB-T2, d) 34DCA-T2, e) 23DCA-T2, f) Heat-killed control (KC). | Excel |
| Table S6 | Absolute abundance of archaeal ASVs, calculated based on the Illumina sequencing data and on qPCR: a) 34DCNB-T2, b) 25DCNB-T2, c) 23DCNB-T2, d) 34DCA-T2, e) 23DCA-T2, f) Heat-killed control (KC). | Excel |

| | | |
|---|---|---|
| Table S7 | Amendments and HPLC results for the bottles Abiotic Control 1 (AC1), Abiotic Control 2 (AC2), and Heat-killed control (KC). | Excel |
| Table S8 | Amendments and HPLC results for the first transfers (T1). | Excel |
| Table S9 | Amendments, HPLC and GC results for the second transfers (T2). | Excel |
| Table S10 | Anion concentrations (by IC) for the second transfers (T2) and AC2. | Excel |
| Table S11 | Chloride balance and sulfate content in enrichment cultures transfer 2 (T2). | Excel |
| Table S12 | Electron balance calculations for the second transfers (T2). | Excel |
| Table S13 | Calculations for yields of *Dehalobacter* during growth on dichloroaniline (this study). | Excel |
| Table S14 | Published *Dehalobacter* cell yields, including substrates. | Excel |

**Text S1: Original microcosm study setup and results.**

The original microcosm study prepared from material from the site in the northeast of Brazil is described in detail in a Master's thesis,[1] but the overview is summarized here. Four different sets of microcosms (1A, 1B, 2A and 2B) were prepared. Set 1A (shallow – 5.2 to 5.8 m depth) and 1B (deep – 5.8 to 6.2 m depth) were constructed using subsurface solid material from well N083 and groundwater from PM19. Set 2A (shallow – 4.8 to 5.4 m depth) and 2B (deep – 5.4 to 6.0 m depth) were constructed from subsurface solid material from well N082 and groundwater from PM12. A total of 96 microcosms (36 aerobic and 60 anaerobic) were constructed in the study, but only the anaerobic microcosms are described here. Site groundwater and solid material were placed inside a disposable anaerobic glove bag (Aldrich-AtmosBagTM) together with additional materials required to construct the various treatment and control microcosms. The glove bag was purged with a carbon dioxide/nitrogen (20:80 %) gas mixture to create an anaerobic environment. The solid material was homogenized to improve reproducibility between replicates and to ensure that control and treatment microcosms contained similar starting materials. Microcosms were constructed by filling sterile 250 mL (nominal volume) glass bottles with 20 milliliters (mL) of homogenized subsurface material and 100 mL of groundwater. The bottles were capped with Mininert™ closures to allow repetitive sampling of the bottle with minimal losses of volatile organic compounds (VOCs), and to allow for amendment of target compounds and nutrients as needed throughout the incubation period. All controls and treatments were constructed in triplicate, and one microcosm in each triplicate group received 150 μL of a 1g/L stock of resazurin as redox indicator.

Sterile control microcosms (SC) were constructed to quantify potential abiotic and experimental losses of target compounds from the microcosms. These controls were constructed as described above but were amended with 1.5 mL of 5% mercuric chloride (equal to a final liquid concentration of 0.05%) and 0.6 mL of 5% sodium azide (equal to a final liquid concentration of 0.02%) to inhibit microbial activity. The SCs made with site material (these did not have added FeS) did not show any obvious signs of nitro-reduction. Unamended active control microcosms (UAC) were constructed to evaluate the ability of indigenous bacteria in the site solid material and groundwater to intrinsically degrade the target aromatic compounds. No electron donor or other amendments were added to these intrinsic controls. Three additional sets of treatments were prepared to test different electron donors and acceptors: 1) donor amended microcosms received soluble electron donors such as ethanol and sodium lactate to a concentration of approximately 100 mg/L each; 2) sulfate amended microcosms received sodium sulfate to a concentration of 2 mM, and 3) nitrate amended microcosms; received sodium nitrate to make a concentration of 2 mM. Microcosms were amended with donor every 2-3 months, and with sulfate and nitrate as needed (sulfate and nitrate concentrations were monitored regularly).

Following construction, all anaerobic microcosms were transferred from the temporary glovebag to an anaerobic glove box (Coy Lab Products) for incubation. During incubation, all microcosms were covered to minimize photodegradation and inverted to prevent VOC losses via the microcosm closure. Groundwater samples were collected from the various control and treatment microcosms every 3 weeks for analysis of aromatics, VOCs and headspace gases (e.g., ethene and methane). Inorganic anions (chloride, nitrate, nitrite, sulfate and phosphate) were monitored approximately every 2-3 months (more frequently at the start of the study). Samples for DNA extraction and subsequent qPCR and sequencing analysis were collected from selected bottles at various times during the first year of monitoring. The microcosm setup, including amendments, compound concentrations and a summary of results, is provided in excel Table S1 and Figure S2. Sequencing results from the subsurface material and the microcosms that were used as inoculum for the anaerobic enrichment cultures described in this study are shown in excel Table S3a-c. The relationship between subsurface material, original microcosms and transfers, and their microbial community is illustrated in Figure 4.

**Text S2: Amendments to enrichment cultures and controls.**

The first set of 5 enrichment bottles (referred to as T1 for transfer #1 from original microcosms) were amended with single compounds as illustrated in Figure S1. Stock solutions of individual dichloronitrobenzenes or dichloroanilines were prepared gravimetrically in acetone. To amend cultures with individual compounds without adding acetone, the following steps were taken. Inside a fumehood, the required volume of stock solution was first dispensed into a sterile 2 mL glass vial (Agilent) and the acetone was evaporated to dryness The vial containing substrate coated on the glass was brought into the glovebox and dropped into the appropriate enrichment bottle. For example, if the desired starting concentration was 10 mg/L 2,3-DCA in 100 mL of culture, then 200 µL of a 5g/L 2,3-DCA stock solution in acetone was evaporated. To remove volatile compounds from the headspace before opening the bottle caps for refeeding, bottles were removed from the glovebox and the headspace was flushed with $N_2$ with a needle through the mininert cap with the cap slightly cracked open. The cap was tightened, needle removed, and the sealed bottle was returned to the glovebox.

The amount of donor added to the bottles was around 17 times, in electron equivalents, of that required for reduction of DCNB to Aniline (nitroreduction and complete dichlorination). For example, for a 10 mg/L feeding concentration, corresponding to 62 µM of DCA or 52 µM of DCNB (considering 10 eeq/mol), 510 µM of ethanol (12 eeq/mol) and 210 µM of lactate (12 eeq/mol) were amended at each feeding of chlorinated acceptor, corresponding to 3 µl of neat ethanol and 33 µl of 0.7 M lactate (or 60% by mass).

Neat ethanol and sodium lactate (60% by mass) were brought into the glovebox after being purged with $N_2$. All bottles, caps, and feeding vials were kept inside the glovebox for at least 3 days before being used.

**Text S3: Details of analytical procedures (pH, HPLC, IC, GC).**

*pH measurements*

A 22G needle was attached to a 1 mL Luer-LokTM tip syringe (BD) to withdraw 1 mL sample from the bottle for pH test. After collection, the sample was transferred into a 1.5 mL polypropylene microcentrifuge tube (Fisher Scientific Co.) and the pH was measured by using a pH Spear meter (Oakton Instruments). Prior to measuring samples, the pH meter was calibrated according to the manufacture's protocol, using pH 4, pH 7, and pH 10 OrionTM buffer solutions (Thermo Scientific).

*High-performance liquid chromatography (HPLC)*

Aniline, chloroanilines (2-, 3-, and 4-CA), dichloroanilines (2,3-, 2,5-, and 3,4-DCA), and dichloronitrobenzenes (3,4-, 2,5-, and 2,3-DCNB) (Sigma Aldrich) were analyzed using a Hewlett-Packard/Agilent 1050 series high performance liquid chromatograph (HPLC) system with a quaternary pump and an autosampler (Agilent Technologies, Santa Clara, CA, USA). The HPLC was equipped with an Acclaim™ 120 C18 column, 3 µm particle size, 4.6 x 150 mm, with average pore diameter of 120 Å and an AcclaimTM C18 guard cartridge, with 5 µm particle size, 4.6 x 10 mm (both from Thermo Scientific). The UV detector was set to 254 nm, mobile phase was 50% Mili-Q water and 50% acetonitrile (Milipore Sigma) at a flow rate of 1 mL/min, isocratic flow.

Samples (2 mL) were collected inside the glovebox using a 22G needle (BD) attached to a Luer-Lock 2 mL gastight glass syringe and taken out of the glove box. In the fume hood, the needle was discarded, and an ethanol pre-washed 0.22 µm Chromspec PTFE Syringe filter 13 mm (Chromatographic Specialties Inc.) was attached to the syringe and a new 22G needle was attached. A portion of the sample (1 mL) was used to flush the filter and was discarded. The remaining sample was filtered into a clear glass 350 µL flat bottom 6x31 mm insert (Chromatographic Specialties Inc.) placed inside a clear 2 mL autosampler glass vial (Agilent Technologies) until it was full. The vial was closed with a PTFE silicone-coated cap (VWR). The HPLC run time was 25 min per sample and the retention times are listed below. Standard concentrations used for calibrations of DCNBs, DCAs, CAs, and aniline, all in methanol (Sigma Aldrich) matrix, were 0.1, 0.5, 1, 2, 5, and 10 mg/L.

Compounds quantified by HPLC and their retention time.

| Compound | Retention time (min) |
|---|---|
| Aniline | 3.3 |
| 4-CA | 4.9 |
| 3-CA | 5.3 |
| 2-CA | 5.4 |

| | |
|---|---|
| 3,4-DCA | 7.8 |
| 2,3-DCA | 9.0 |
| 2,5-DCA | 10.0 |
| 2,3- and 2,5-DCNB (co-eluted) | 14.0 |
| 3,4-DCNB | 16.5 |

*Ion chromatography (IC)*

Anions, including lactate, acetate, propionate, formate, butyrate, pyruvate, chloride, nitrate, nitrite, sulfate and phosphate, were measured by ion chromatography (IC) using a Dionex™ Integrion IC system with a Dionex™ IonPac™ AS11-HC analytical column. The eluent was potassium hydroxide (KOH) generated automatically by the eluent generator at a flow of 0.25 mL/min. A gradient elution was used starting at 0.5 mM KOH increasing to 2.5 mM in 10 mins and further increasing to 30 mM by 29 mins. The concentration was held at 30 mM for another 11 mins before returning to the starting conditions of 0.5 mM by 42.1 mins and held steady for 4.9 mins for re-equilibration. External standards were prepared in water with concentrations ranging from 0.005 to 1 mM. Samples (~1 mL) were collected from culture and control bottles with a disposable syringe and were filtered through 0.2 μm nylon filters (Fisher Scientific) prior to injection on the IC. The retention time for each analyzed anion is shown below. Standard concentrations used for calibrations were 0.005, 0.01, 0.05, 0.2, 0.5, and 1 mM. The total run time was 47 min per sample, and the retention times are listed below.

Anions quantified by IC and their retention time.

| **Anion** | **Retention time (min)** |
|---|---|
| Lactate | 11.8 |
| Acetate | 12.2 |
| Propionate | 13.3 |
| Formate | 14.4 |
| Butyrate | 15.0 |
| Pyruvate | 15.8 |
| Chloride | 19.4 |
| Nitrite | 20.9 |
| Nitrate | 25.3 |
| Sulfate | 28.3 |
| Phosphate | 35.4 |

*Gas chromatography (GC)*

These analyses were performed more frequently during the microcosm experiment for volatile DCB isomers. Dichlorobenzenes (1,2-, 1,3-, 1,4-DCB) (Sigma Aldrich) and methane (Scott) were used as standards and analyzed using the Agilent 7890A gas chromatograph (GC) with headspace autosampler G1888, equipped with a GSQ-Plot column (0.53 mm x 30 m) (both from Agilent Technologies) and a flame ionization detector (FID). The temperature of the packed inlet was set to 200°C, the detector was set to 250°C, and the carrier gas during the run was helium. $H_2$ flow was 40 mL/min, air flow was 400 mL/min, and total flow was 11 mL/min. The oven was programed as follows: 35°C for 1.5 min, ramp 15°C/min to 100°C, ramp 5°C/min to 185°C hold 10 min, ramp 20°C/min to 200°C, hold 10 min. The total run time was 43.6 min per sample vial and the retention times are listed below.

Compounds quantified by GC and their retention times.

| Compound | Retention time (min) |
|---|---|
| Methane | 0.9 |
| Benzene | 15.2 |
| 1,3-DCB | 32.6 |
| 1,4-DCB | 33.4 |
| 1,2-DCB | 34.4 |

The autosampler setting was as follows: oven at 70°C; loop at 80°C; transfer line at 90°C; vial equilibration time 40 min; pressurization time: 0 min; loop fill time: 0.2 min; loop equilibration time: 0 min; inject time: 3 min; GC cycle time: 47 min; shaking: low.

Liquid samples were collected using a 22G (0.7 mm x 40 mm) PrecisionGlideTM needle (BD) attached to a Luer-Lock 2 mL gastight glass syringe. In the fume hood, clear glass flat bottom 10 mL autosampler vials (Agilent Technologies) were filled with 5 mL of acidified water (2.4 mL of HCl 5 N topped up to 1 L with Mili-QTM water). The needle was placed into the acidified water, the sample was rapidly dispensed, and the vial was crimped with an open top 20 mm aluminum crimp seal with PTFE/silicone coated septum (200 mm, 130mil, white) (both from Chromatographic Specialties Inc.) by using a vial crimping tool.

**Text S4: Quantitative polymerase chain reaction (qPCR).**

The abundance of total bacterial and total archaeal 16S rRNA genes was measured using a CFX96TM real-time PCR detection system (qPCR) with a C1000 thermocycler (Bio-Rad Laboratories Inc). The 20 µl qPCR reactions were prepared in a PCR cabinet (ESCO Technologies,) and were made up by 10 µl of SsoFastTM EvaGreen® SuperMix (Bio-Rad Laboratories Inc.), 1 µl of each forward and reverse primers (10 uM stock, making final concentration of 250 nM for both primers), 6 µl of UV treated UltraPure Distilled water (Invitrogen), and 2 µl of DNA extract diluted 10x. The qPCR cycle was as follows: 98°C for 2 min, 40 cycles of 98°C for 5 seconds, annealing at 55°C (total bacteria) or 60 °C (total archaea) for 10 seconds, followed by an increase from 65°C to 95°C at 0.5°C increments over 10 seconds. Calibration curves for qPCR were prepared from serial dilutions of target-containing

*Dehalococcoides* (for general bacteria) and *Methanomethylovorans* (for general archaea) plasmids with concentrations ranging between $10^1$ and $10^8$ gene copies/μL (prepared by doing serial dilutions with IDTE buffer (10 mM Tris, 0.1 mM EDTA, pH 8, Integrated DNA Technologies IDT), to avoid adsorption to the tubes). A no-template control was also included in each run. All qPCR samples were run in triplicate. The final concentration (copies/mL) was calculated by multiplying the total mean starting quantity values from qPCR (copies/μL), the dilution factor, and the elution volume during DNA extraction (μL), divided by the amount of culture filtered (mL). The $R^2$, slope, reaction efficiency, y-intercept and values for standards and no template controls for each qPCR plate run are provided below.

**Text S5: Preparation of Bubble Plots and Non-metric multidimensional Scaling (NMDS) analysis.**

Bubble plots showing absolute abundance of different ASVs as copies/mL were made in R, using the ggplot2[2] and reshape2 libraries.

Non-metric multidimensional scaling (NMDS) plots were created in order to visualize overall grouping patterns of ASVs for the various cultures. The sample matrix was subsampled to an even depth of 29000 reads in all samples. Out of the 46 sequenced samples collected up until day 520, one sample had quite few reads (<2000 reads) and was therefore excluded from the NMDS analysis. NMDS plots were constructed using the "ordinate" function within the phyloseq package v1.34.0[3] in R v4.0.1, which leverages functions from the vegan package v2.5-7.[4] The NMDS was based on a Bray-Curtis dissimilarity matrix with two dimensions (k = 2). The plot was created using the "plot_ordination" function, which is the general ordination plotter in phyloseq and is based on ggplot2.

**Text S6: 16S rRNA gene sequences of *Dehalobacter* ASV1 (detected in 34DCNB-T2) and *Dehalobacter* ASV2 (detected in 23DCA-T2 and 34DCA-T2) obtained from 16S Illumina sequencing.**

*Dehalobacter* ASV1:
GGGCCCGCACAAGCGGTGGAGCATGTGGTTTAATTCGACGCAACGCGAAGAACCTTA
CCAAGGCTTGACATCCAACTAATCCCGTAGAGATATGGGAGTGCCCTTCGGGGAAAGT
TGAGACAGGTGGTGCATGGTTGTCGTCAGCTCGTGTCGTGAGATGTTGGGTTAAGTCC
CGCAACGAGCGCAACCCCTATATTTAGTTGCTAACAGGTAAAGCTGAGAACTCTAGAT
AGACTGCCGGTGACAAACCGGAGGAAGGTGGGGATGACGTCAAATCATCATGCCCCT
TATGTCTTGGGCTACACACGTGCTACAATGGACGGTACAGACGGAAGCGAAGCCGCG
AGGTGAAGCAAATCCGAGAAAGCCGTTCTCAGTTCGGATTGCAGGCTGCAACTCGCCT
GCATGAAGTCGGAATCGCTAGTAATCGCAGGTCAGCACACTGCGGTGAATACGTTCCC
GGGCCTT

*Dehalobacter* ASV2:

GGGCCCGCACAAGCGGTGGAGCATGTGGTTTAATTCGACGCAACGCGAAGAACCTTA
CCAAGGCTTGACATCTACAGAATCCTTAAGAGATTAGGGAGTGCCTTTCGGGGAACTG
TAAGACAGGTGGTGCATGGTTGTCGTCAGCTCGTGTCGTGAGATGTTGGGTTAAGTCC
CGCAACGAGCGCAACCCCTATATTTAGTTGCTAACAGGTAAAGCTGAGAACTCTAGAT
AGACTGCCGGTGACAAACCGGAGGAAGGTGGGGATGACGTCAAATCATCATGCCCCT
TATGTCTTGGGCTACACACGTGCTACAATGGACGGTACAGACGGAAGCGAAGCCGCG
AGGTGAAGCAAATCCGAGAAAGCCGTTCTCAGTTCGGATTGCAGGCTGCAACTCGCCT
GCATGAAGTCGGAATCGCTAGTAATCGCAGGTCAGCACACTGCGGTGAATACGTTCCC
GGGCCTT.

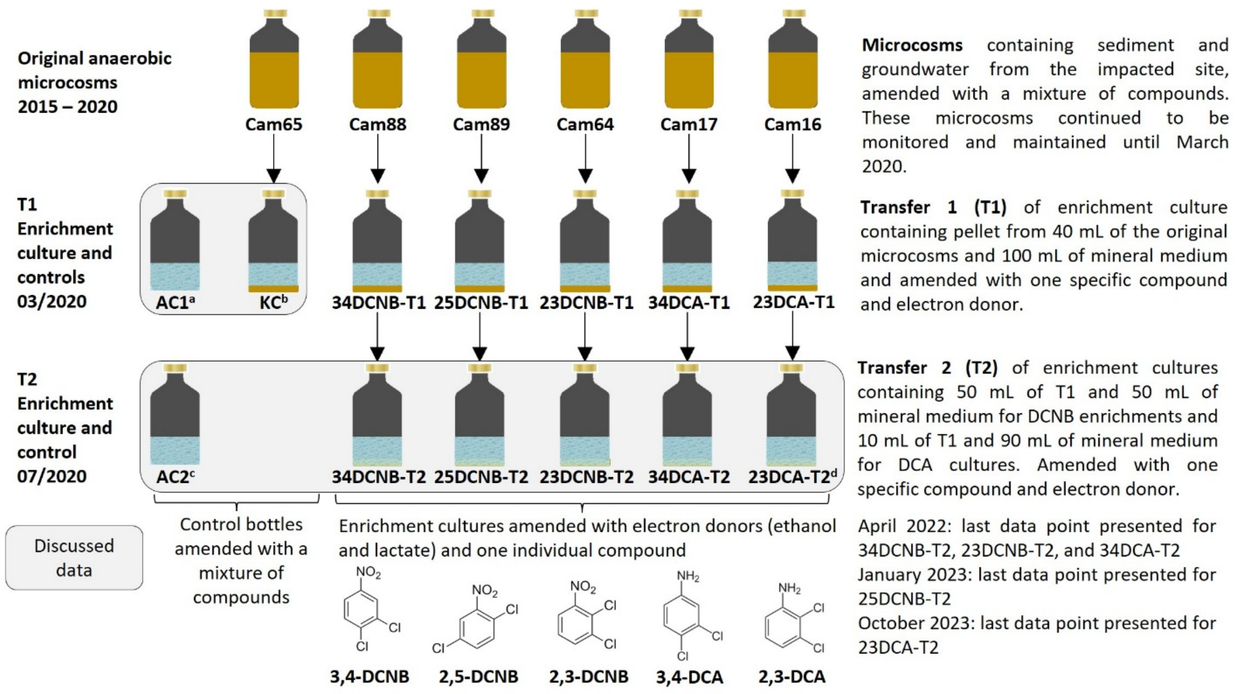


**Figure S1. History of enrichment cultures, showing setup details of original microcosms, T1, and T2 transfers, including controls.** Data from T2 transfers are the focus herein.

Footnotes:

[a]Abiotic control #1 (AC1) contained mineral medium, 3,4-DCNB, 2,3-DCNB, 3,4-DCA, 2,3-DCA, 2-CA, aniline, and donors.

[b]The heat killed-control (KC) contained an autoclaved cell pellet from original microcosms, mineral medium 3,4-DCNB, 2,5-DCNB, 3,4-DCA, 2,3-DCA, 2-CA, aniline, and donors.

[c]Abiotic control #2 (AC2) contained mineral medium free of FeS and with 3,4-DCNB, 2,3-DCNB, 3,4-DCA, and 2,3-DCA.

[d]A subsample of 23DCA-T2 was taken on day 653 (April 21st, 2022) for additional experiments and enrichment (data in companion manuscript).[5]

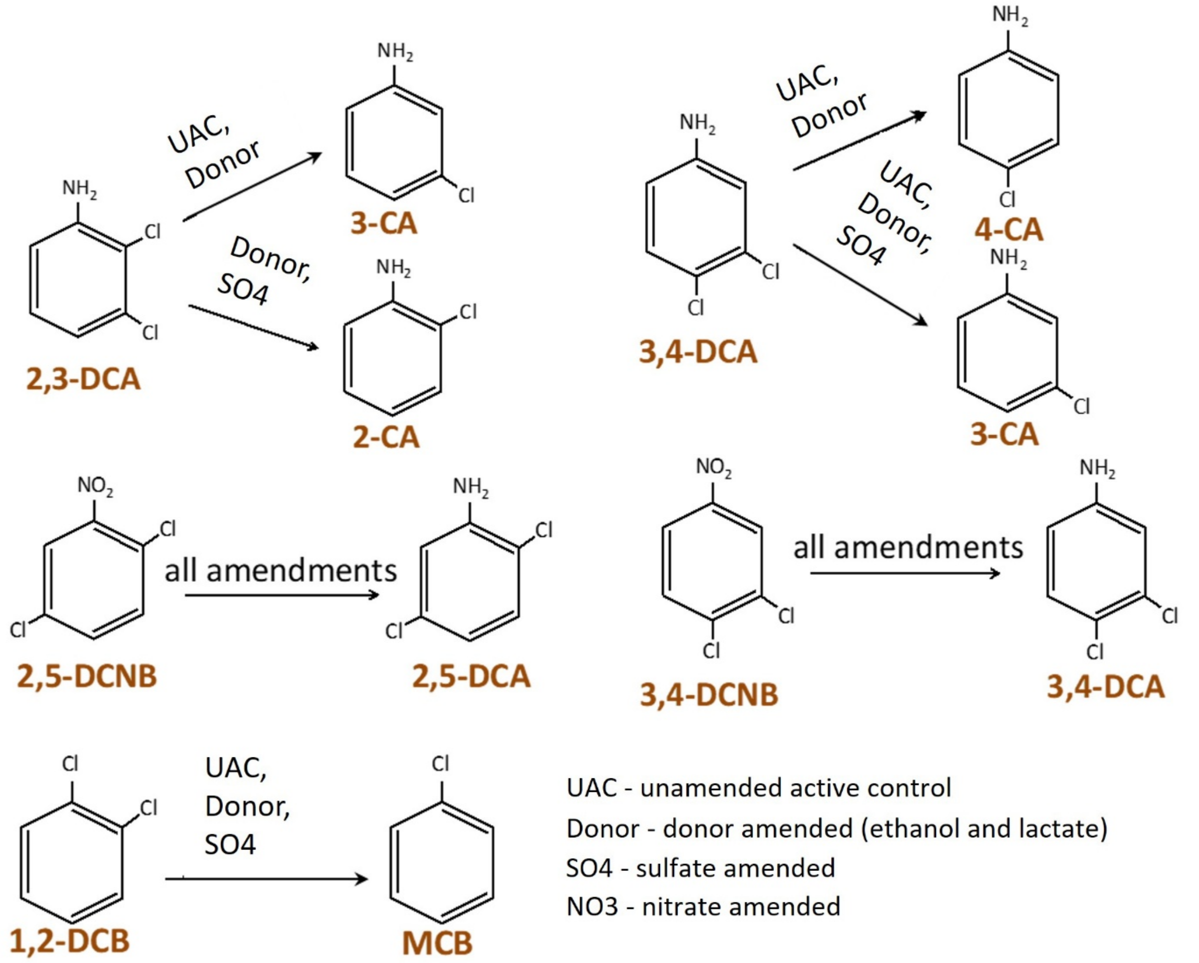


**Figure S2. Overview of observed transformations deduced in active anaerobic microcosms.**

Amendments/conditions are indicated on arrows. The sterile control microcosms (SC) made with site material (these did not have added FeS) did not show any obvious signs of nitro-reduction.

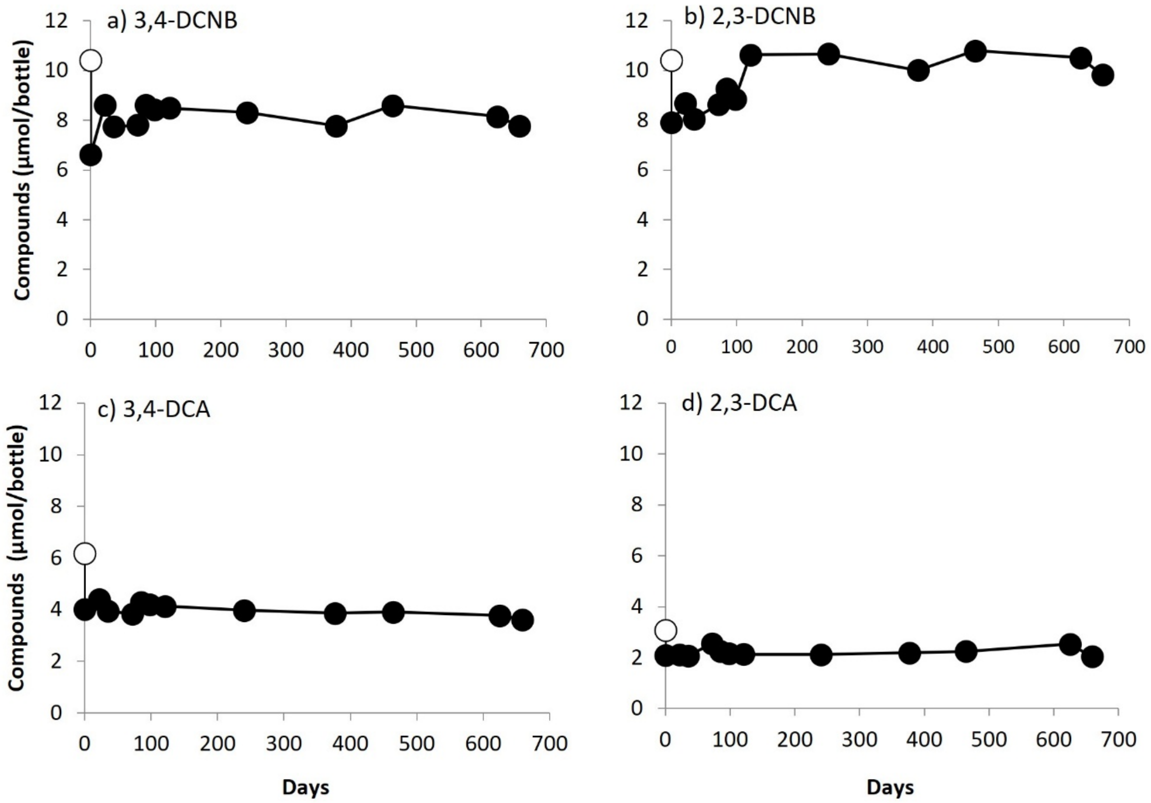


**Figure S3. Concentration profiles in control bottle AC2, medium with no FeS.** Concentration of 3,4-DCNB (a), 2,3-DCNB (b), 3,4-DCA (c), and 2,3-DCA (d) over time in the abiotic control containing mineral medium free of FeS (AC2). **NOTE: all compounds were in the same bottle.** Open white symbols are amounts fed to the bottle at T=0. No transformation observed of any compound.

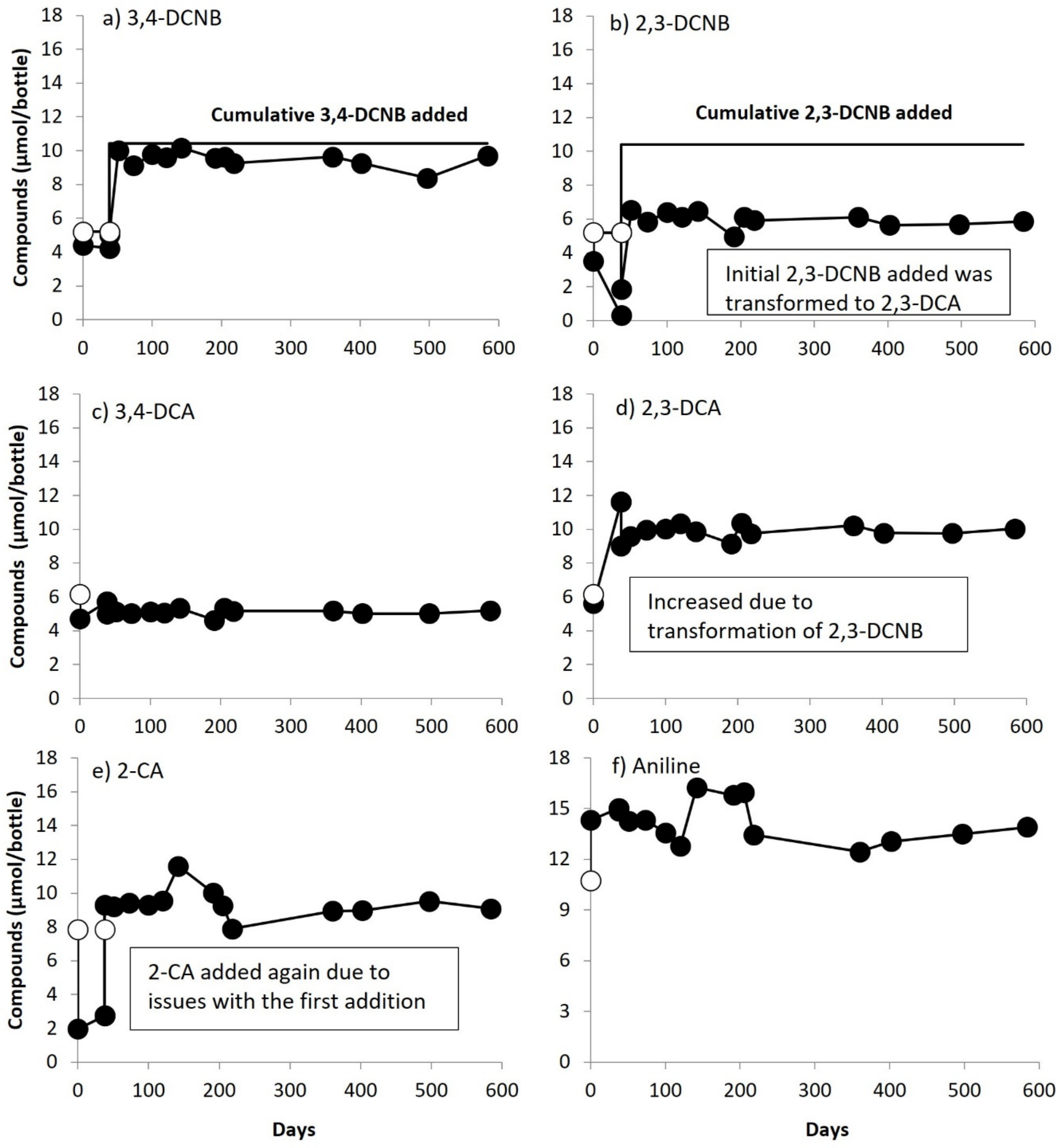


**Figure S4. Concentration profiles in control bottle AC1, medium with FeS.**
Concentrations of 3,4-DCNB (a), 2,3-DCNB (b), 3,4-DCA (c), 2,3-DCA (d), 2-CA (e), and aniline (f) over time in the abiotic control bottle containing mineral medium with FeS (AC1). **NOTE: all compounds were in the same bottle.** Open white symbols are amounts fed to the bottle at T=0 and at T=50 for 3,4-DCNB and 2,3-DCNB. **One feeding of 2,3-DCNB was reduced, but subsequently concentration remained stable. 3,4-DCNB was not reduced**.

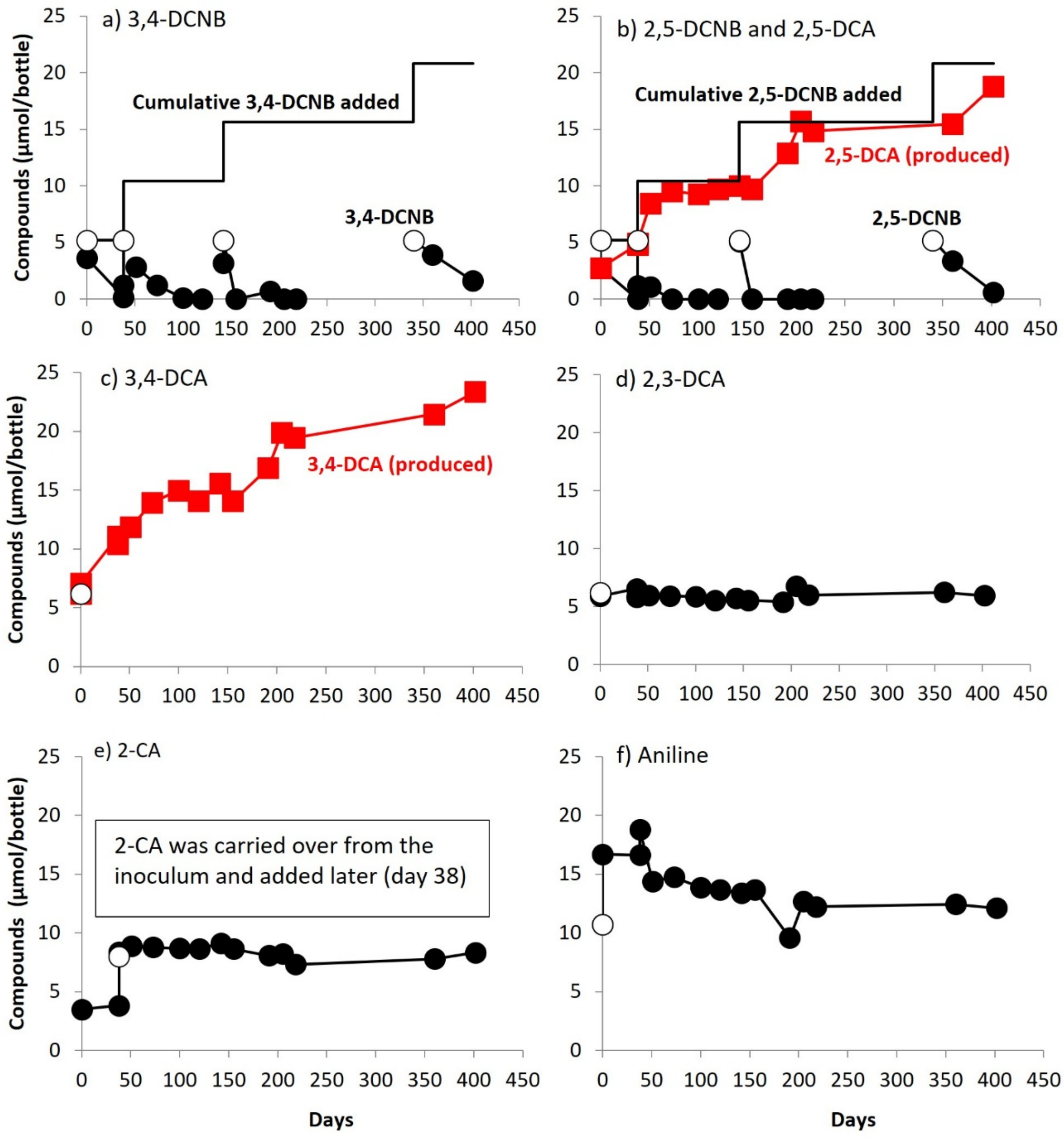


**Figure S5. Concentration profiles in autoclaved (killed) control bottle KC.** Concentration of 3,4-DCNB (a), 2,5-DCNB and 2,5-DCA (b), 3,4-DCA (c), 2,3-DCA (d), 2-CA (e), and aniline (f) over time in the heat-killed control (KC). **NOTE: all compounds were in the same bottle**. Open white symbols are amounts fed to the bottle at T=0 and at T=38, T=142, and T=340, for 3,4-DCNB and 2,5-DCNB. **3,4-DCNB and 2,5-DCNB were repeatedly reduced with production of corresponding dichloroanilines.**

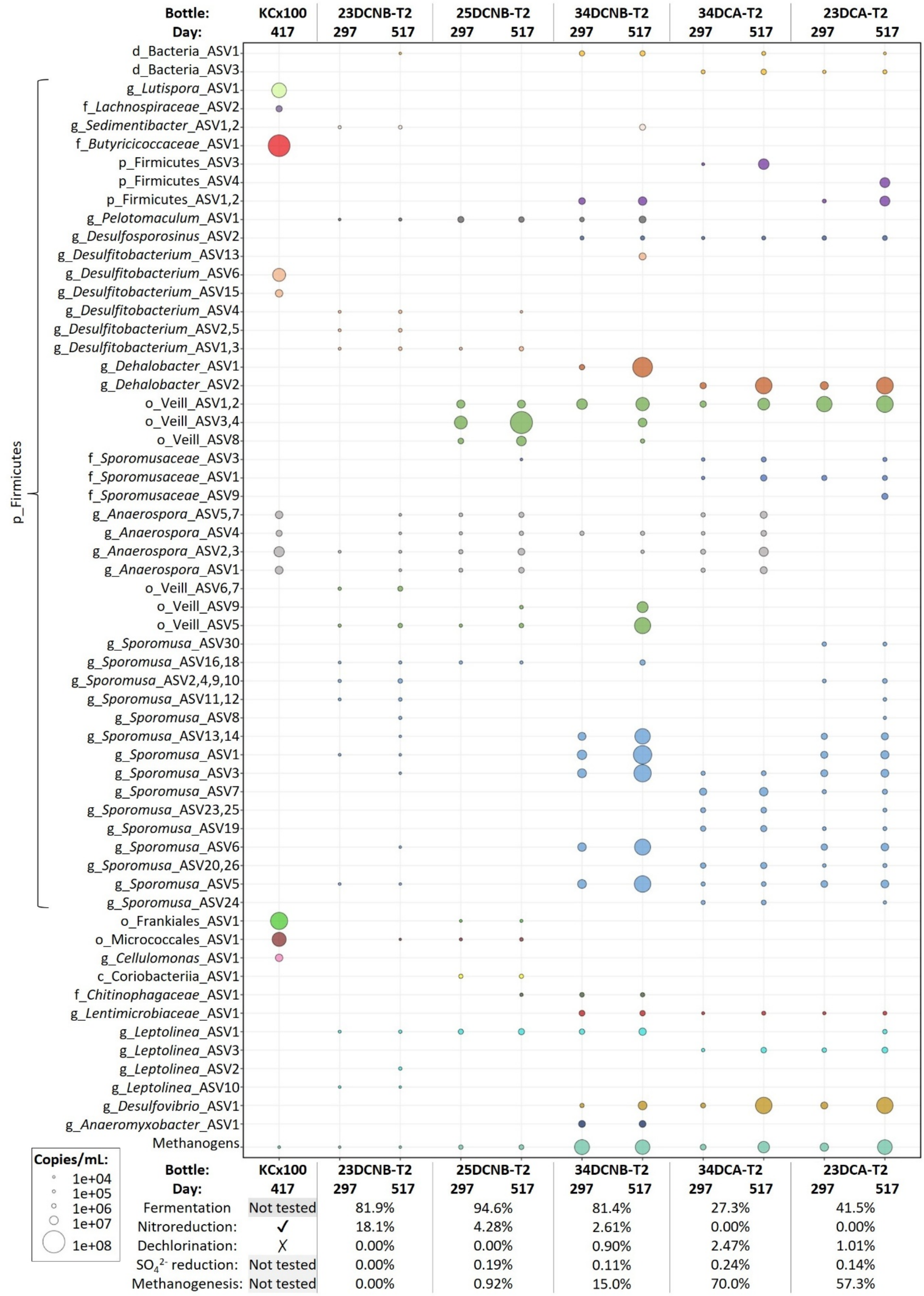
Bottle: KCx100 23DCNB-T2 25DCNB-T2 34DCNB-T2 34DCA-T2 23DCA-T2
Day: 417 297 517 297 517 297 517 297 517 297 517
d_Bacteria_ASV1
d_Bacteria_ASV3
g_Lutispora_ASV1
f_Lachnospiraceae_ASV2
g_Sedimentibacter_ASV1,2
f_Butyricicoccaceae_ASV1
p_Firmicutes_ASV3
p_Firmicutes_ASV4
p_Firmicutes_ASV1,2
g_Pelotomaculum_ASV1
g_Desulfosporosinus_ASV2
g_Desulfitobacterium_ASV13
g_Desulfitobacterium_ASV6
g_Desulfitobacterium_ASV15
g_Desulfitobacterium_ASV4
g_Desulfitobacterium_ASV2,5
g_Desulfitobacterium_ASV1,3
g_Dehalobacter_ASV1
g_Dehalobacter_ASV2
o_Veill_ASV1,2
o_Veill_ASV3,4
o_Veill_ASV8
f_Sporomusaceae_ASV3
f_Sporomusaceae_ASV1
f_Sporomusaceae_ASV9
g_Anaerospora_ASV5,7
g_Anaerospora_ASV4
g_Anaerospora_ASV2,3
g_Anaerospora_ASV1
o_Veill_ASV6,7
o_Veill_ASV9
o_Veill_ASV5
g_Sporomusa_ASV30
g_Sporomusa_ASV16,18
g_Sporomusa_ASV2,4,9,10
g_Sporomusa_ASV11,12
g_Sporomusa_ASV8
g_Sporomusa_ASV13,14
g_Sporomusa_ASV1
g_Sporomusa_ASV3
g_Sporomusa_ASV7
g_Sporomusa_ASV23,25
g_Sporomusa_ASV19
g_Sporomusa_ASV6
g_Sporomusa_ASV20,26
g_Sporomusa_ASV5
g_Sporomusa_ASV24
p_Firmicutes
o_Frankiales_ASV1
o_Micrococcales_ASV1
g_Cellulomonas_ASV1
c_Coriobacteriia_ASV1
f_Chitinophagaceae_ASV1
g_Lentimicrobiaceae_ASV1
g_Leptolinea_ASV1
g_Leptolinea_ASV3
g_Leptolinea_ASV2
g_Leptolinea_ASV10
g_Desulfovibrio_ASV1
g_Anaeromyxobacter_ASV1
Methanogens
Copies/mL:
1e+04
1e+05
1e+06
1e+07
1e+08
Bottle: KCx100 23DCNB-T2 25DCNB-T2 34DCNB-T2 34DCA-T2 23DCA-T2
Day: 417 297 517 297 517 297 517 297 517 297 517
Fermentation Not tested 81.9% 94.6% 81.4% 27.3% 41.5%
Nitroreduction: ✓ 18.1% 4.28% 2.61% 0.00% 0.00%
Dechlorination: ✗ 0.00% 0.00% 0.90% 2.47% 1.01%
SO4 2- reduction: Not tested 0.00% 0.19% 0.11% 0.24% 0.14%
Methanogenesis: Not tested 0.00% 0.92% 15.0% 70.0% 57.3%

**Figure S6. [above – previous page] Overview of the most abundant bacterial ASVs in the active bottles (ASVs > 1%, 2 time points), and activities observed in bottles.**

Veillonellales-Selenomonadales order is shortened to “Veill”. The table below the graph represents electron accepting processes (% electron equivalents) in each culture. For the “killed” control KC, we did not measure ions and methane.

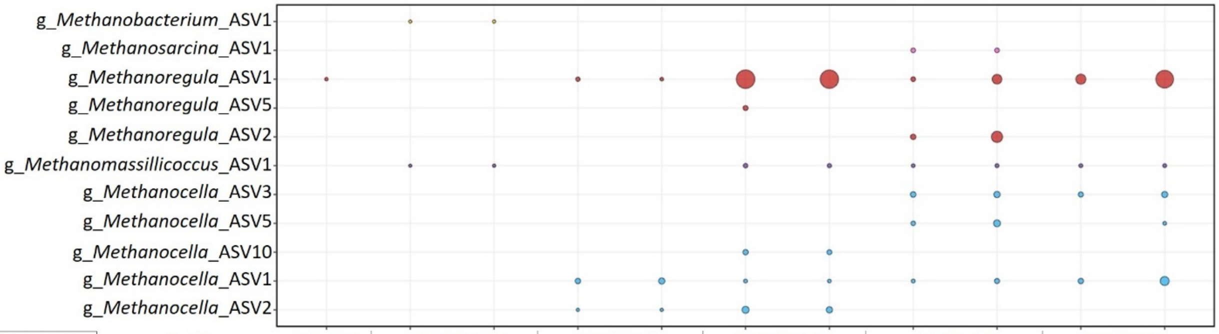


Copies/mL: 1e+04, 1e+05, 1e+06, 1e+07, 1e+08

| Bottle: | KCx100 | 23DCNB-T2 | 25DCNB-T2 | 34DCNB-T2 | 34DCA-T2 | 23DCA-T2 |
|---|---|---|---|---|---|---|
| Day: | 417 | 297 517 | 297 517 | 297 517 | 297 517 | 297 517 |
| Fermentation | Not tested | 81.9% | 94.6% | 81.4% | 27.3% | 41.5% |
| Nitroreduction: | ✓ | 18.1% | 4.28% | 2.61% | 0.00% | 0.00% |
| Dechlorination: | ✗ | 0.00% | 0.00% | 0.90% | 2.47% | 1.01% |
| $SO_4^{2-}$ reduction: | Not tested | 0.00% | 0.19% | 0.11% | 0.24% | 0.14% |
| Methanogenesis: | Not tested | 0.00% | 0.92% | 15.0% | 70.0% | 57.3% |

**Figure S7. Overview of the most abundant archaeal ASVs in the active bottles (ASVs > 1%, 2 time points), and activities observed in the bottles.**

The table below the graph represents electron accepting processes (% electron equivalents) in each culture. For the “killed” control KC, we did not measure ions and methane.

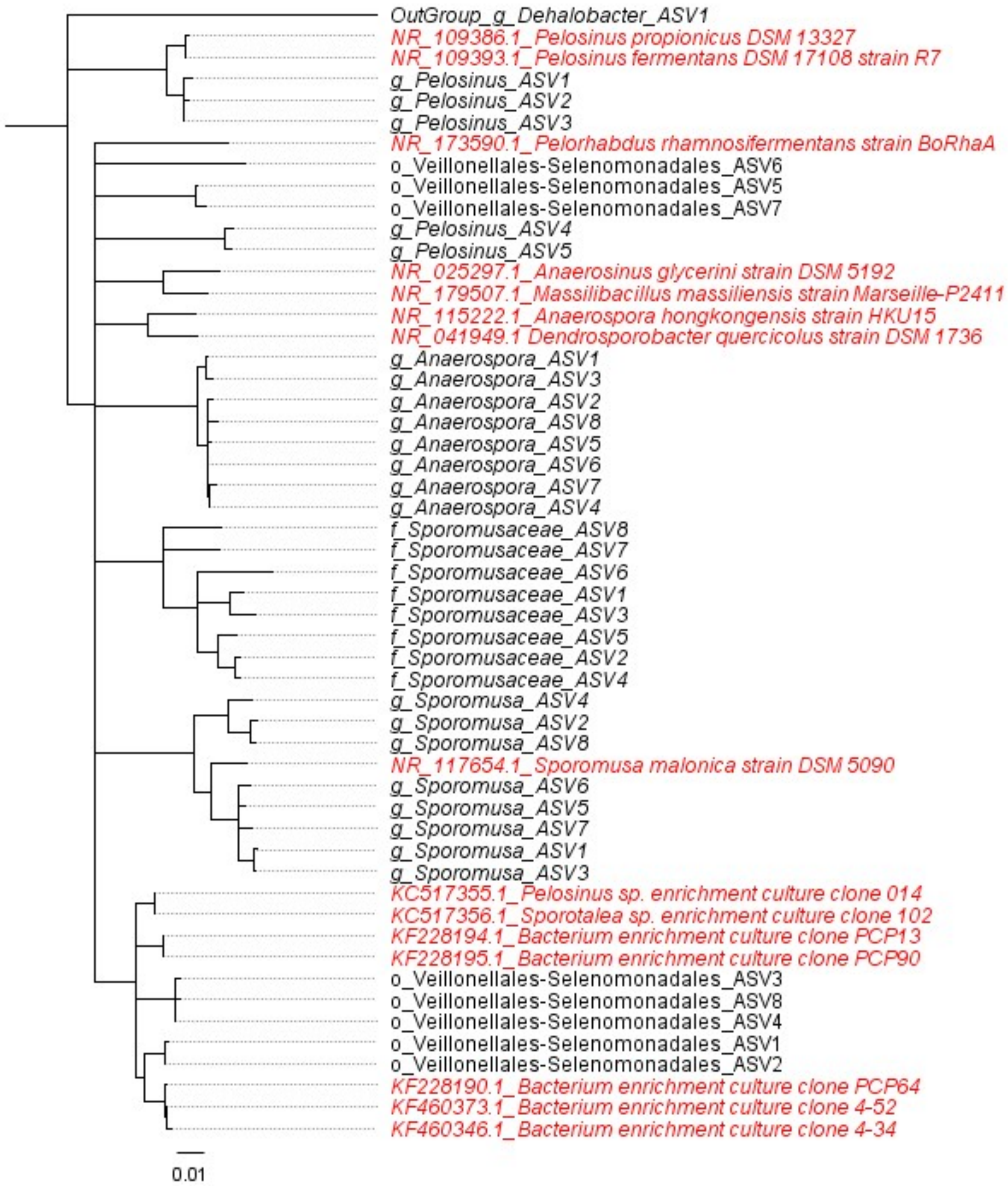


**Figure S8. Phylogenetic tree of organisms classified as part of the Veillonellales-Selenomonadales order.**

Organisms in black are the most abundant Veillonellales-Selenomonadales ASVs identified in the enrichment cultures of this study: 8 classified at the order level Veillonellales-Selenomonadales (among them, 6 were >1% in at least one sample from 25DCNB-T2: ASV1 to 5 and ASV8); 8 further classified at family level *Sporomusaceae*; and at the genus level, 8 classified to *Sporomusa*, 8 to *Anaerospora*, and 3 to *Pelosinus*. The *Dehalobacter* ASV1 detected in 34DCNB-T2 culture was included as an outgroup, also shown in black. The 15 organisms in red are the closest matches to the Veillonellales-Selenomonadales_ASV1 (most abundant ASV across the cultures) obtained by running the Basic Local Alignment Search Tool (BLAST) available at the National Center for Biotechnology Information (NCBI) when the parameters "Highly similar sequences (megablast)" or "Somewhat similar sequences (blastn)" were used for the search. All the sequences were aligned and truncated using the function MUSCLE alignment; the tree was built using Geneious Tree Builder (Global alignment with free and gaps, 65% similarity, Tamura-Nei, Neighbor-Joining) on Geneious 8.1.9. The units of branch length represent nucleotide substitutions per site.

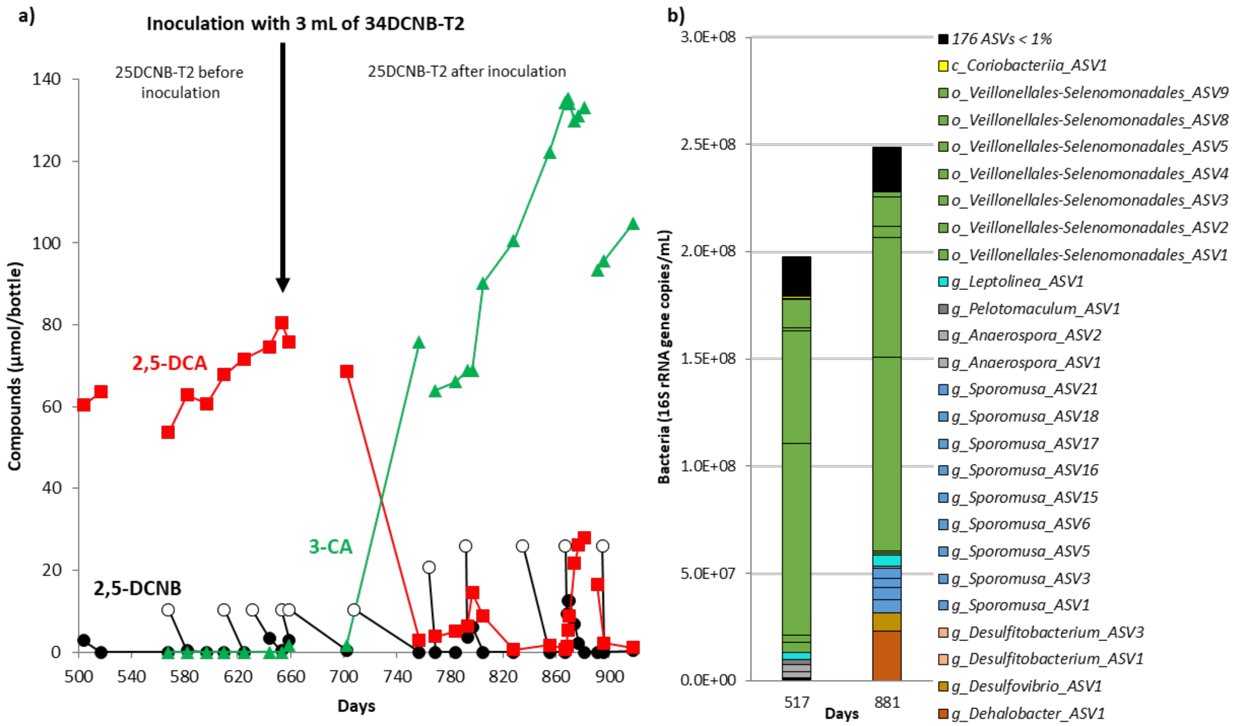


**Figure S9. Activity in culture 25DCNB-T2 between days 500 and 920, before and after the culture was inoculated with 3 mL of 34DCNB-T2 on Day T=653 (a).**

2,5-DCA that previously accumulated in the culture was reduced after inoculation, forming 3-CA. NOTE: Mineral medium was added to the culture at T=528, T=659, T=764, and T=881, and thus the compounds were diluted. Open white symbols are amounts of 2,5-DCNB fed to the bottle on specific days. **DNA samples collected from 25DCNB-T at T=517 (before inoculation) and T=881 show growth of *Dehalobacter*_ASV1 (same ASV detected in 34DCNB-T2 went from 0 to ~9% of the bacterial community) (b) (Data provided in excel Table S5b).**

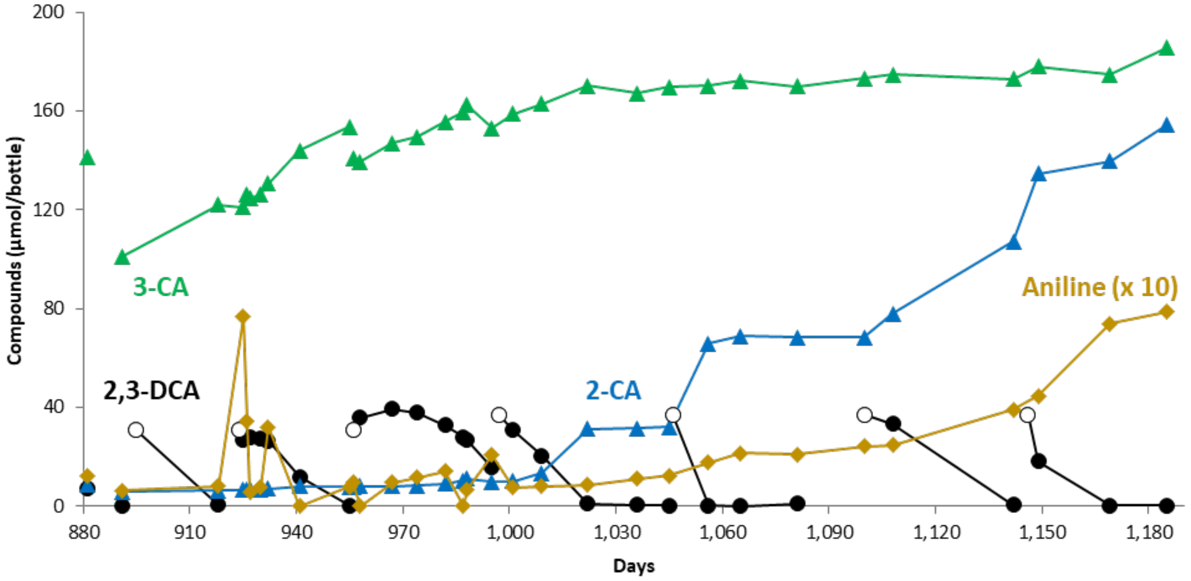


**Figure S10. Shift in activity in culture 23DCA-T2: 2-CA production and progressive accumulation of aniline.**

Concentrations of aniline are multiplied by 10. Mineral medium was added to the culture at T=881 and T=956, and thus the compounds were diluted. Open white symbols are amounts of 2,3-DCA fed to the bottle at multiple time points.

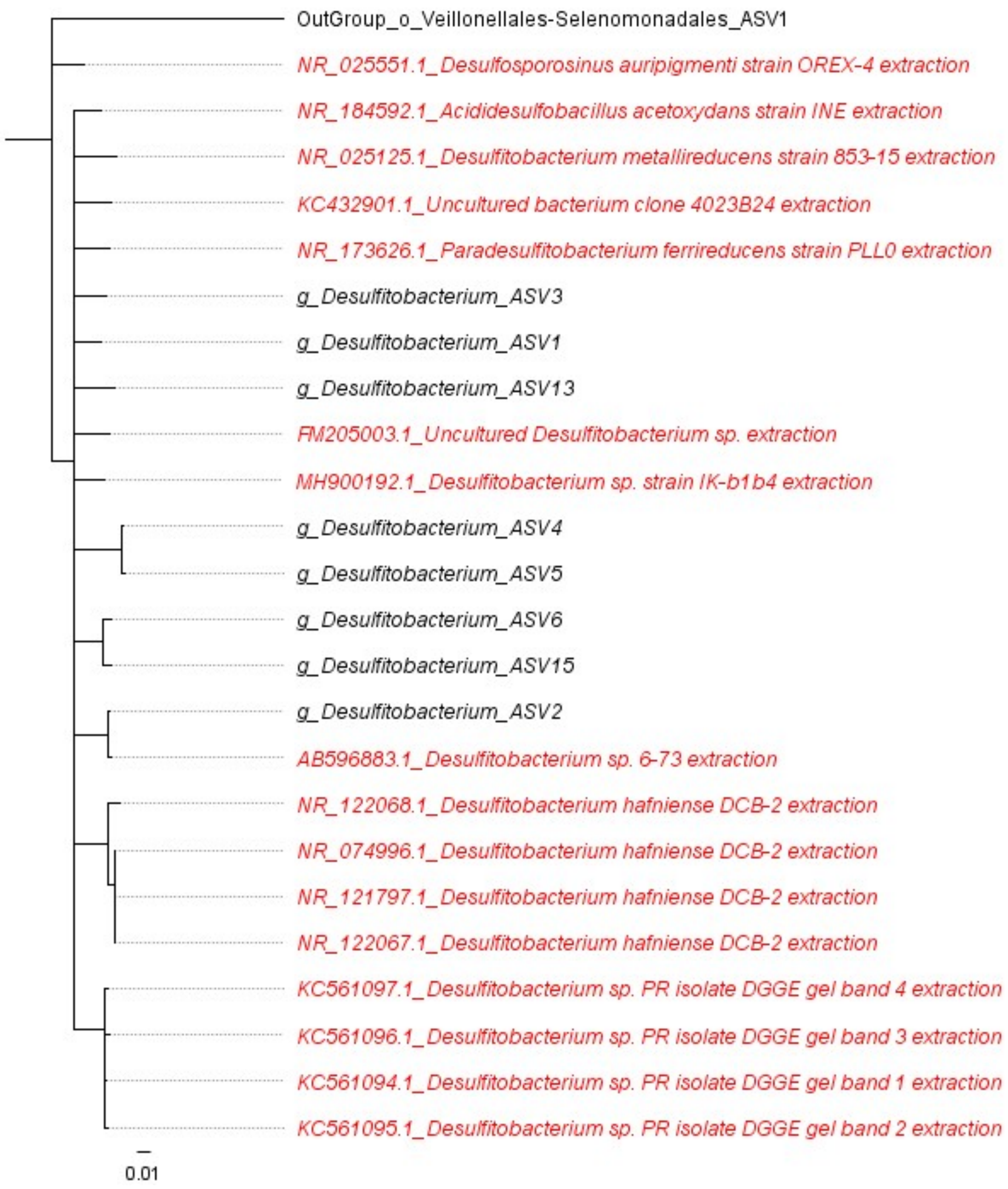


**Figure S11. Phylogenetic tree of organisms classified as part of the *Desulfitobacterium* genus.**

Organisms in black are the most abundant *Desulfitobacterium* ASVs identified in the enrichment cultures of this study. The Veillonellales-Selenomonadales ASV1 detected in 34DCNB-T2 culture was included as an outgroup, also shown in black. The 16 organisms in red are the closest matches to the *Desulfitobacterium*_ASV1 (most abundant ASV across the cultures) obtained by running the Basic Local Alignment Search Tool (BLAST) available at the

National Center for Biotechnology Information (NCBI) when the parameters “Highly similar sequences (megablast)” or “Somewhat similar sequences (blastn)” were used for the search. All the sequences were aligned and truncated using the function MUSCLE alignment; the tree was built using Geneious Tree Builder (Global alignment with free and gaps, 65% similarity, Tamura-Nei, Neighbor-Joining) on Geneious 8.1.9. The units of branch length represent nucleotide substitutions per site.
